# Shear Stress Modulates Inflammatory Responses in Porcine Endothelial Cells Contributing to the Fibrotic Response of Porcine Cardiac Fibroblasts

**DOI:** 10.1101/2025.02.03.636305

**Authors:** Pengfei Ji, Kathryn Jane Grande-Allen, Swathi Balaji, Ravi K. Birla, Sundeep G. Keswani

## Abstract

This study is focused on discrete subaortic stenosis (DSS) which is a congenital heart disease resulting in the formation of fibrotic tissue in the left ventricular outflow track of pediatric patients [1]. The pathogenesis of the molecular mechanism resulting in the formation of the fibrotic tissue remains unknown and is the focus of this study. We hypothesized the cytokine released in response to wall shear stress act on the fibroblasts to promote the formation of the fibrotic membrane and subsequently, DSS. To test this study hypothesis, porcine endocardial endothelial cells were cultured within cone and plate bioreactors, designed to replicate the wall shear stress observed in the left ventricular outflow track of DSS patients. Conditioned media was collected as a function of shear stress magnitude and used to condition porcine cardiac fibroblasts. We used an extensive set of end-point metrics to characterize fibroblast phenotype to include bulk-RNA sequencing, RT-qPCR, cell viability and immunofluorescence. The results of this study demonstrate that shear stress-conditioned media from porcine EEC releases a defined cocktail of cytokines that provide the trigger to program the fibrotic response during DSS. These results provide specific molecular targets that can be developed into a therapeutic strategy for patients with DSS and provide a solution to an otherwise challenging disease to manage.

## INTRODUCTION

DSS is a well-known cardiovascular disease and results in the isolated obstruction of left ventricular outflow tract (LVOT) in pediatric patients [2]. Central to the understanding of DSS is an understanding of the surgical methodologies, the molecular mechanism governing recurrence and changes in the aortic and mitral valve function resulting from the formation of the fibrotic tissue [3]. The study of DSS study involves changes in the hemodynamics environment of the LVOT, subsequent changes in the wall shear stress on the endothelial cells and fibrotic response of the underlying fibroblasts. Despite several reports or literature on DSS, none of them have figured out the answer to these questions entirely.

DSS manifests as a distinct circular fibromuscular rim of tissue, featuring a variable-width fibrous inner ring [4]. In addition, DSS can form or present other types or morphologies except fibromuscular ring of tissue, as it can also manifest as an incomplete shelf and ridge-like structure [5]. It has been reported that there are five tissue layers related to DSS including, endothelial layers, sub-endothelial layer with glycosaminoglycans, fibroelastic layer containing collagen bundles and elastin fibrils, smooth muscle layer with a thickened basement membrane, and fibrous layer exhibiting heightened collagen content [6].

The mechanisms and formation of DSS have been disputed and the basis of disagreement for a long time. Even though DSS was believed to be a congenital heart disorder, there was evidence and reports that DSS was an acquired lesion [3]. Currently, the therapeutic method for DSS is surgical resection. However, DSS has a high risk of recurrence even after operation, notably, DSS frequently appears pristine and gives the impression of no previous resection during reoperation [7]. In a recent study, a more extensive myocardial resection was not recommended as the patient did not decrease the reoperation rate and complete alleviation of the obstruction and posed a high risk of heart block [2].

Furthermore, the shear stress of force on the endothelial cell layer of the LVOT has been reported and resulted in cell proliferation and induced cellular growth and division [8, 9]. The postulated mechanism stated that the shear stress impaired or injured the endocardial layer, and the cellular wound attracted the fibroblasts to repair the damage. Also, cell injury released inflammatory cytokines that stimulate the fibroblast to differentiate into contractile and collagen-secreting myofibroblasts. Persistent myofibroblast proliferation may result in the additional accumulation of a fibromuscular layer, producing DSS in the LVOT [10]. The flow disturbances that lead to DSS are often minuscule and after successful resection, identifying the root cause is typically not evident upon examination through the aortic valve, and the fibrotic tissue may reappear after a few years even though achieving a complete operative relief [11].

While the hemodynamic etiology of DSS is widely acknowledged, a comprehensive validation of this conception is firmly established. Among the almost 400 publications on DSS, the predominant focus on echocardiographic delineations of LVOT hemodynamics, exploration of genetic inheritance, and biochemical characterizations of DSS lesions. Recognizing that the pathogenesis of DSS may arise from a confluence of hemodynamic and genetic factors prompt the exploration of investigated approaches [12]. Therefore, mechanobiological experiments were designed to simulate the unidirectional flow of blood between the left ventricle and the aorta. This method operates in mechanically challenging environments, subjecting the system to various stresses, including bending and tensile stress, pressure, and fluid shear stress [13].

The cone-and -plate bioreactor (CPB) is the device that generates a nearly consistent shear stress in the culture environment. This device comprises an inverted cone that rotates above a flat stationary plate, and fluid circulates in the gap between the cone and the plate [14]. The regulation of the shear stress distribution on the plate surface is based on adjusting the angular velocity applied to the rotating cone. There are a lot of advantages of the CPB that make this device a good choice for subjecting cells to pulsatile or oscillatory shear stresses defined by high shear stress rates. The flow produced in this device has undergone comprehensive investigation through experiments [15, 16, 17]. Based on the above discussion, the CPB generates a flow aligned along the direction and a consistent shear stress on the bottom of the plate surface defined as T=pvw[r/(h+ra)] [18, 19].

In the context of the DSS project, the CPB will be used to generate the uniform shear stresses on EECs as the study observed fluid shear stresses in the LVOT of DSS patients. The magnetic stir plate regulates the angular velocity of the cone by adjusting the RPM on the control unit. From our previous work (unpublished work), 4 different shear stress regimes were designed for this study, negative control, positive control, low shear stress, and high shear stress, based on the published computational models for DSS. Negative control = 0 dynes/cm^2^ or 0 RPM, Positive control = 6 dynes/cm^2^ or 100 RPM, low shear stress = 15 dynes/cm2 or 280 RPM and high shear stress = 35 dynes/cm2 or 660 RPM [12, 20, 21].

Alternatively, the DSS in LV exhibited a steep aortoseptal angel that supported mechanobiological etiology, which suggests that hemodynamic abnormalities in the LV interacted with the endocardium to drive fibrosis and the formation of DSS membranes. Additionally, the increment of the shear stress magnitude and gradient captured on the septal wall conducted on fibrosis and explain the frequent association between DSS and steep LVOTs [11, 12, 22]. The current finding indicates the presence of a crucial region on the septal wall where shear stress alternations are maximized. This region may also correspond to the development of fibrosis and the formation of DSS lesions [21]. Thus, further studies are necessary to confirm the sensitivity of EECs to the shear stress features of the LVOT, and the formation of fibrous membrane tissues in the LVOT.

Given the above, cardiac fibrosis develops following heart injury, inflammation, or aging, leading to the accumulation of extracellular matrix (ECM) and declination of the cardiac function. The primary cell type implicated in the fibrotic remodeling process is the cardiac fibroblast based on their roles in ECM production. It is important to understand cardiac fibroblast, and how to mitigate the detrimental effects of fibroblast activation. There remains considerable uncertainty regarding cardiac fibroblasts’ origin, function, gene expression and signaling pathways [23, 24]. Therefore, this study aimed to elucidate the role of porcine cardiac fibroblasts in the development and progression of DSS. The cardiac fibroblasts were estimated and revealed the roles of pathological progression and potential involvement in metabolic pathways associated with the DSS. This research proposed that developmental mechanisms can play important roles in promoting the development of cardiovascular disorders as well as cardiac repair and regeneration.

## MATERIALS AND METHODS

### Porcine EECs and Fibroblast Culture

A 0.1% gelatin solution (Gelatin methacryloyl, bloom 300, degree of substitution 60%, Sigma-Aldrich, 900622-1G) was used to coat the bottom of the cell plates and incubated for 15 minutes. The gelatin coating was used to promote attachment and proliferation of both porcine EECs and fibroblasts. Frozen vials were retrieved from liquid nitrogen for both porcine EECs and fibroblasts and thawed by placing the vials in a 37℃ water bath until the contents were 70% thawed. Subsequently, fibroblast growth medium 3 (PromoCell Inc., Catalog # C-23010) was used to culture the fibroblasts and endothelial cell medium (Sciencell, Catalog # 1001) was used for the culture of porcine EECs, both according to manufacturers instructions. Porcine EECs were cultured and expanded in 10 cm plate and then transferred to 60 mm plates at a density of 500,000 cells per plate and cultured for an additional 2 days, after which the porcine EECs reached a confluency of 70%. At this point, the porcine EECs were ready for bioreactor studies. Similarly, fibroblasts were plated at an initial density of 100,000 cells per well in 6 well plates and allowed to expand for 2 days until reaching a confluency of 50% prior to the media transfer studies.

### Conditioning medium treatment with Porcine fibroblasts

Porcine EECs were subjected to bioreactor conditioning at 0, 6, 15, and 35 dyne/cm^2^ for 24 hours, and the conditioned medium was collected for all 4 different bioreactor treatments. The conditioning medium and fibroblast medium were prepared in 3 different ratios including 10%, 25%, and 50%. Subsequently, the conditioned medium was used to treat the porcine fibroblasts for 48 hours, and a monitoring system recorded the porcine fibroblast conditions every 24 hours over the timeline of 0, 24, and 48 hours. Based on the results of these studies, a single ratio of conditioned media to fibroblast media was selected and used for all subsequent studies. As a further validation step, the porcine fibroblasts were cultured for 7 days with the conditioned medium at an appropriate ratio from described above.

### Cell culture monitoring systems and Cell Viability Staining

Porcine fibroblasts were monitored by IncuCyte®S3 with conditioning medium treatment for 24, and 48 hours, and following the experiment with a 7-day condition medium treatment at a suitable percentage. The monitor records each time point, to be specific, for the 48 hours of treatment including 0, 24, and 48 hours. For the time course study over the 7 day culture period, time points of day 0, day 4, and day 7 were selected. The IncuCyte®S3 live-cell analysis system from ESSEN Bioscience, Inc., serves as a flexible assay platform that has a cell culture incubator. It automatically captures and analyzes phase and fluorescent images for several days. In this study, the 4x and 10x objectives were used to acquire the images. In addition, porcine fibroblasts were stained with DAPI (Thermo Scientific™ DAPI Solution (1 mg/mL), #EN62248) to assess cell viability.

### RNA extraction, cDNA synthesis, and quantitative real-time PCR

Total RNA was isolated by using the RNeasy Micro Kit (Invitrogen catalog # 12183018A), and the quantity and quality of total RNA were estimated by NanoDrop™ 2000/2000c Spectrophotometers (Thermo Scientific™, #ND2000CLAPTOP). The absorbance of RNA reading at A260/280 within 1.9-2.0 was defined as the acceptable purity range for RNA. Continuously, the total RNA was used to synthesize cDNA using Script™ Reverse Transcription Supermix (Bio-Rad, #1708841). Subsequently, quantitative real-time PCR was performed in duplicate on the CFX384 Touch™ Real-Time PCR Detection System (Bio-Rad, #1845099) using SsoAdvanced™ Universal SYBR® Green Supermix (Bio-Rad, #1725271). The relative mRNA expression levels were normalized with a housekeeping porcine gene GAPDH (Bio-Rad, #10042976).

### Immunofluorescences for Porcine Fibroblasts

The porcine fibroblasts were plated onto 6-well plates and 2 mL of fibroblast growth medium was added per well. Then, the cells were fixed with 4% PFA and the PFA washed with PBS three times. Next, the blocking buffer was prepared which consisted of 3% BSA and 1% Triton-X, and this buffer was added to each well and incubated at room temperature. The porcine fibroblasts were stained with BD Pharmingen™ Alexa Fluor® 488 Mouse Anti-Human Vimentin antibody (BD Biosciences, #562338) with a 1:5 dilution for 1 hour. Subsequently, DAPI was added using a dilution of 1:5000 in each well and stored in the dark for 15 mins. After that, the antibody and DAPI were removed. Finally, the porcine fibroblasts were washed with PBS three times, and images were acquired with a KEYENCE BZ Microscope (BZ-X800).

### RNA sequencing of porcine fibroblasts

The porcine fibroblasts were treated with conditioned media from porcine EECs after bioreactor treatment for 24 hours at 0, 6, 15, and 35 dyne/cm^2^. Genomic analysis was done by Novogene Inc. The workflow of RNA sequencing involved completing a transcriptome for either a single cell or a specific group of cells under defined conditions. Analysis of the transcriptome enables the identification of the differentially expressed genes across distinct cell populations, and can also lead to an understanding of gene boundary identification, variable cleavage, and transcript variation [25]. RNA sequencing utilized Illumina platforms, which operate based on the sequencing-by-synthesis (SBS) mechanism. This technology offers numerous advantages, including high throughput, high accuracy, and low sample requirements. It serves as a powerful tool for investigating RNA transcriptional activity.

### Statistical Analysis

Statistical analyses were performed using SPSS 23.0 software and ImagJ. Data were presented as mean ± SEM. Significance was determined through unpaired two-tailed student t-test, post-hoc Tukey’s testing, one-way analysis of variance (ANOVA), and/or two-way ANOVA. A significance level of P < 0.05 was considered indicative of statistical significance. Additional details can be found in the figure legends.

## RESULTS

### Dose-Response Study – Effect of % Conditioned Media on Fibroblast Culture

To detect the porcine fibroblasts’ response to the porcine EEC conditioned medium, there were four different groups of the conditioned medium as defined here: the normal porcine EECs without treatment were regarded as negative control; the porcine EECs conditioned with 6 dyne/cm^2^ in the CPB were considered as positive control; the low and high shear stress were defined as the 15 dyne/cm^2^ and 35 dyne/cm^2^ in the CPBs. All the porcine EEC medium were from endothelial cell medium. Continuously, the conditioned medium was mixed with porcine fibroblasts medium at 10%, 25%, and 50% for dose titration studies, the porcine fibroblasts were treated with these mixed mediums for 48 hours (2 days). As shown In Fig. 1 (A, B & C), the 10% conditioned media did not result in the cell number decreasing of porcine fibroblasts, and over the 48-hour culture period, the cell number gradually increased due to cell proliferation. In addition, the normal porcine EEC medium (endothelial cell medium) without bioreactor treatment did not exert a detrimental effect on the proliferation and growth of the porcine fibroblasts in the negative control group when compared to other groups. Interestingly, the positive control group did not influence the proliferation of the porcine fibroblasts except the 50% high concentration group. However, in the 50% conditioned media group, the positive control group resulted in a decrease in the cell number, which maybe due to the high concentration of the conditioned medium, which may also include the high concentration of cytokines that caused the decreasing of the cell number of the porcine fibroblasts. From Fig. 1. B, the low and high shear stress groups resulted in decreased cell numbers of the porcine fibroblasts compared to the other group at 0 & 24 hours. However, there was no significant difference among the groups at 48 hours (about 2 days). The most likely reason for this was that the porcine fibroblasts tolerated the conditioned medium during these times. In Fig. 1. C, the positive control, low, and high shear stress groups all resulted in decreased cell numbers of the porcine fibroblasts compared to the other groups during the 48-hour culture time at 50% conditioned medium. The porcine EEC conditioned medium with low and high shear stress results in porcine fibroblast cell death, while the normal porcine EEC medium did not induce a similar reduction in the cell number of porcine fibroblasts. Based on the results of these studies, we concluded the 25% porcine EEC conditioned media was optimal for the culture of porcine fibroblasts and was retained for all future studies.

**Figure 1.**
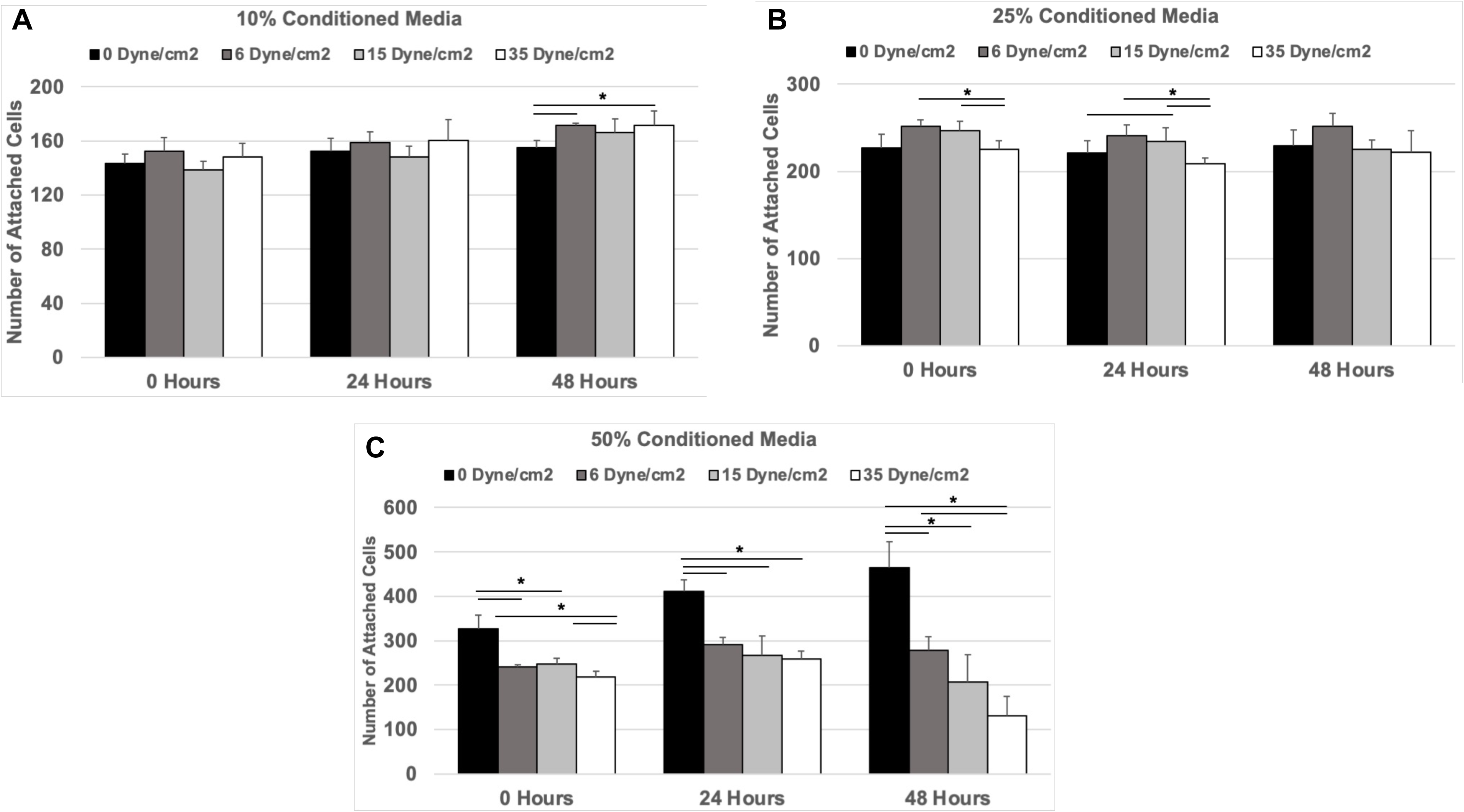
The Effects of % Conditioned Media on Fibroblast Cell Culture. -the porcine fibroblasts were monitored using brightfield images during the time point 0, 24, and 48 hours with 3 different ratios of conditioned medium, 10%, 25%, and 50%. Negative control was defined as porcine fibroblast treatment with porcine EEC conditioning medium (0 dyne/cm^2^ on the CPB). Positive control was defined as porcine fibroblast treatment with porcine EEC-conditioned medium (6 dyne/cm^2^ CBP). Low shear stress was defined as porcine fibroblast treatment with porcine EEC-conditioned medium (15 dyne/cm^2^ CPB). High shear stress was defined as porcine fibroblast treatment with porcine EEC conditioning medium (35 dyne/cm2 on the CPB). A time course study was conducted and cell counts were obtained at the start of the study and 24 and 48 hours after. Brightfield images were obtained using 4x magnification and used to quantify cell counts. We used N=4 independent samples for every time point and all 3 conditioned media formulations (10%, 25%, and 50%), all 4 shear stress conditions (0, 6, 15, and 35 dyne/cm2) at all 3-time points (0, 24 and 48 hours). Results are expressed as mean±SEM. Statistical analysis using two-tailed t test, * P<0.05.

### Time Course Study - Viability of Porcine Fibroblasts as a Function of Time Using 25% Conditioned Media

In this study, the porcine fibroblasts were cultured using 25% conditioned medium treatment since our previous study had shown that 25% is the optimal choice for this study. We used, the Incucyte S3 to monitor the porcine fibroblasts for 7 days, and shown in Fig. 2. A, the cell number of porcine fibroblasts gradually increased before 4 days, but the cell viability of porcine fibroblasts decreased after 4 days, especially day 7. To be more specific, the 15 dyne/cm^2^ and 35 dyne/cm^2^ groups showed reduced cell numbers during the culture time, and there was a significant difference among 6 dyne/cm^2^, 15 dyne/cm^2,^ and 35 dyne/cm^2^ on day 4 and day 7. The DAPI staining of the porcine fibroblasts showed cell confluency with the timeline, and it was observed that day 1 and day 4 increased the cell numbers compared to day 7 in Fig. 2. B. The images were analyzed by ImagJ, and gave several statistical differences in Fig. 2. C. From the DAPI analyzed data, the 0 dyne/cm^2^, 15 dyne/cm2, and 35 dyne/cm^2^ were significantly different among these groups at day 7, especially when the cell number of 35 dyne/cm^2^ decreased.

**Figure 2.**
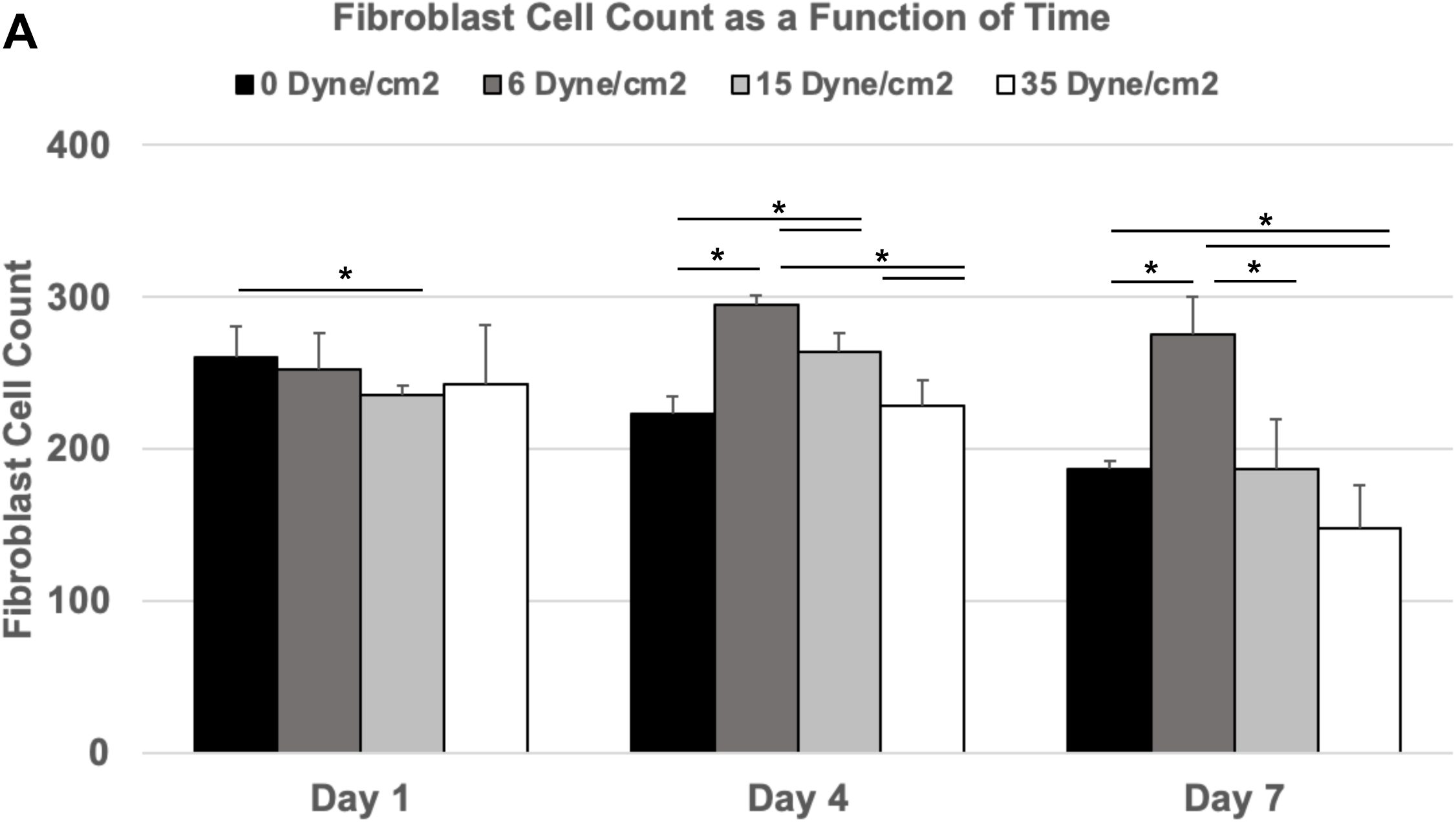
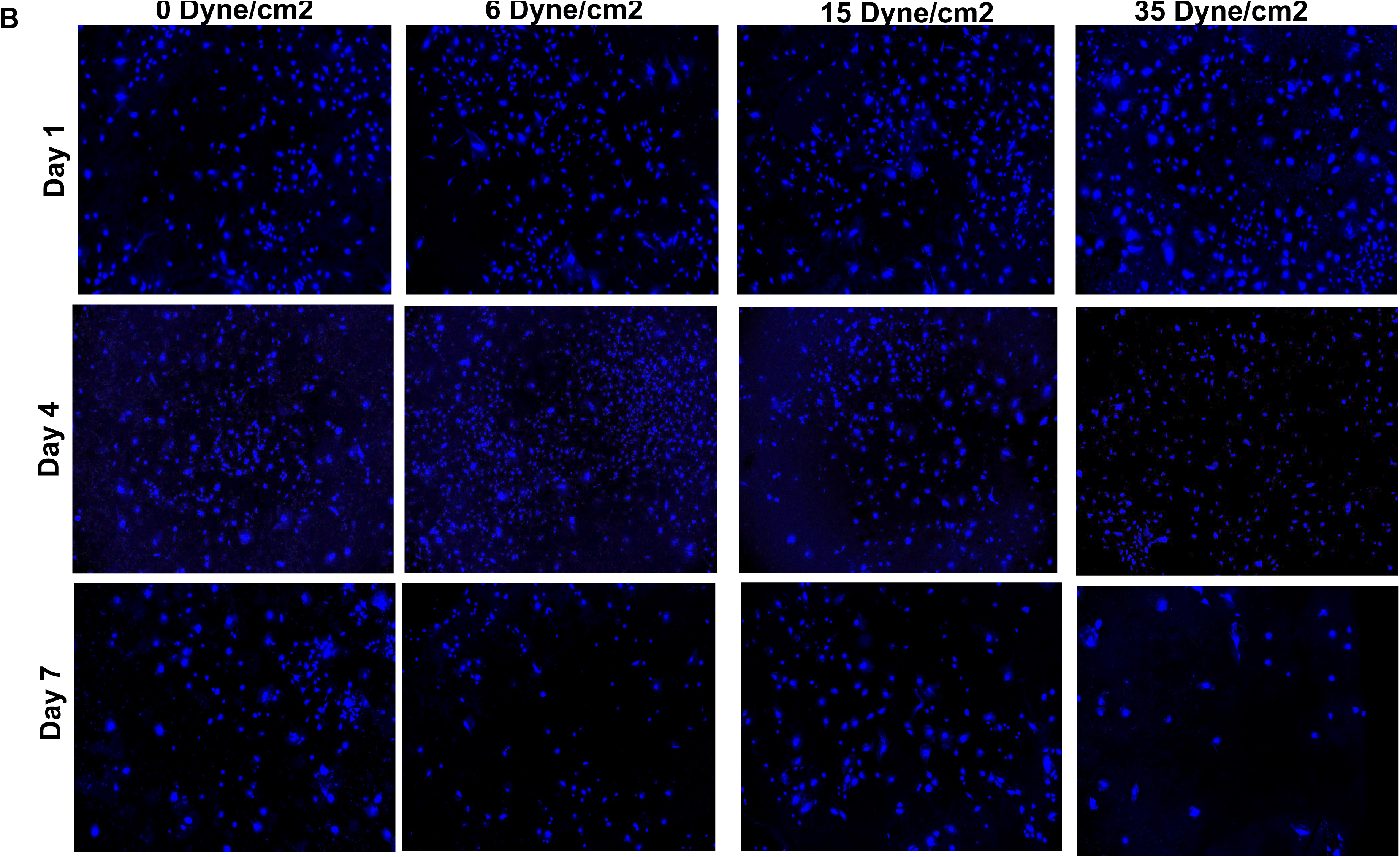
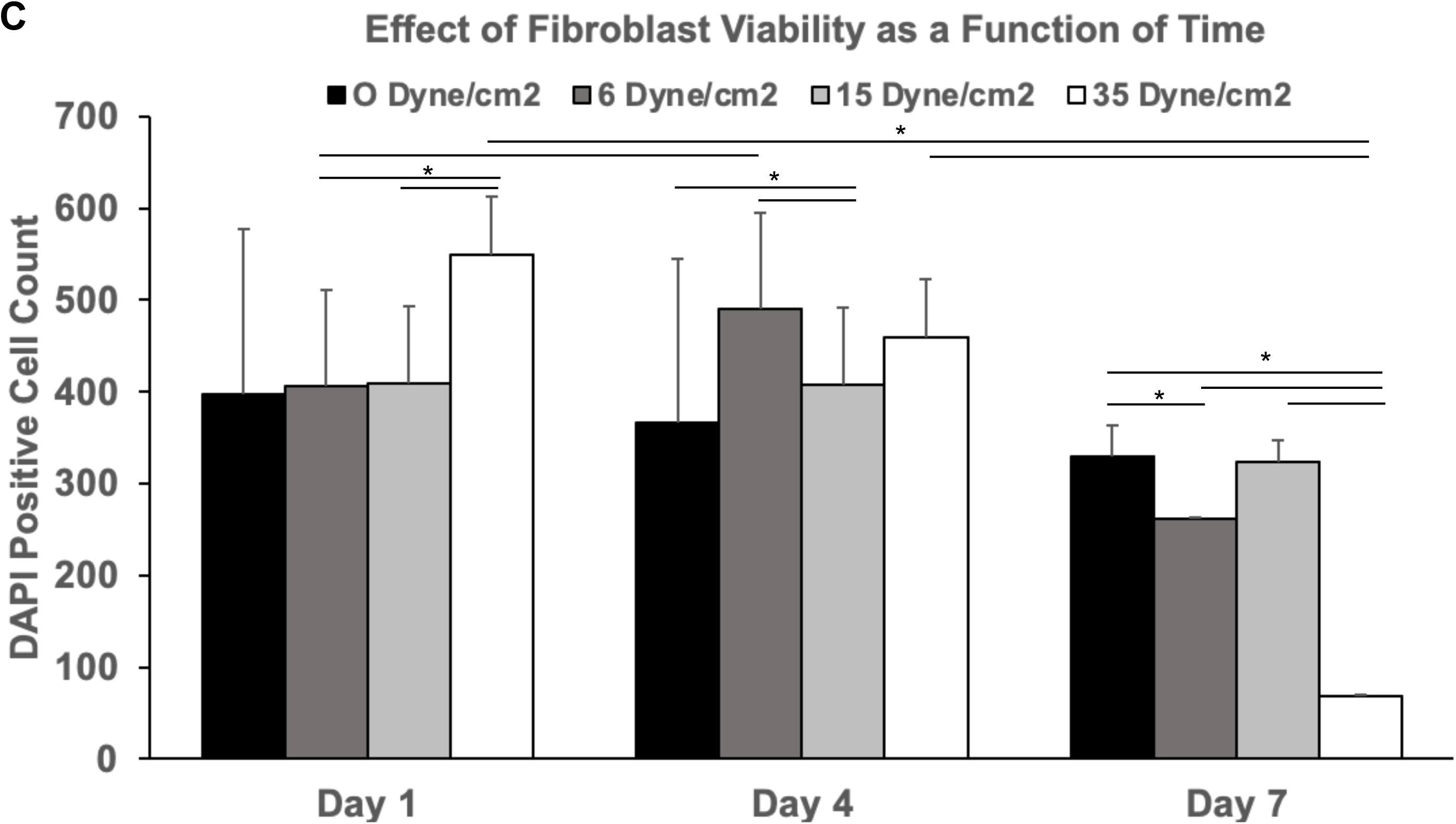
Viability of Porcine Fibroblasts as a Function of Time using 25% Conditioned Media. – Porcine fibroblasts were cultured using 25% and cell viability measured using DAPI staining after day 1, day 4, and day 7. All images are under 4x objectives, and the arrow showed the porcine fibroblasts. Results are expressed as mean±SEM. Statistical analysis using two-tailed t test, * P<0.05.

### Increased Fibrosis in Porcine Fibroblasts Cultured in Conditioned Media

After the shear stress treatment to the porcine fibroblasts, RNA was extracted from porcine fibroblasts on day 1, day 4, and day 7. RT-qPCR was used to reveal the gene expression of the fibrosis markers. In Fig. 3, we found the relative RNA expression of skeletal alpha-actin (ACTA1) was significantly increased in 15 dyne/cm^2^ compared to 6 dyne/cm^2^ at day 7.

**Figure 3.**
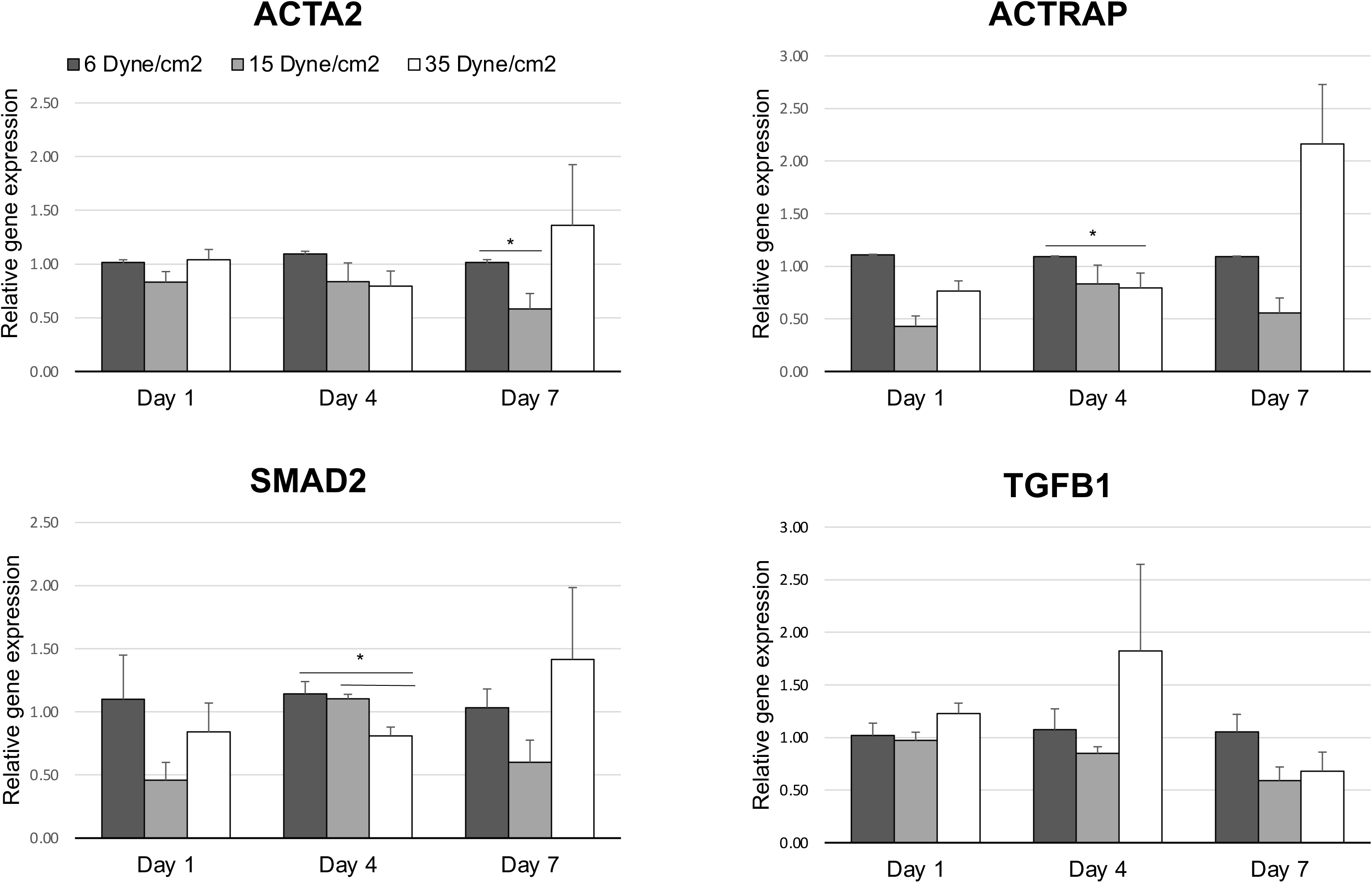
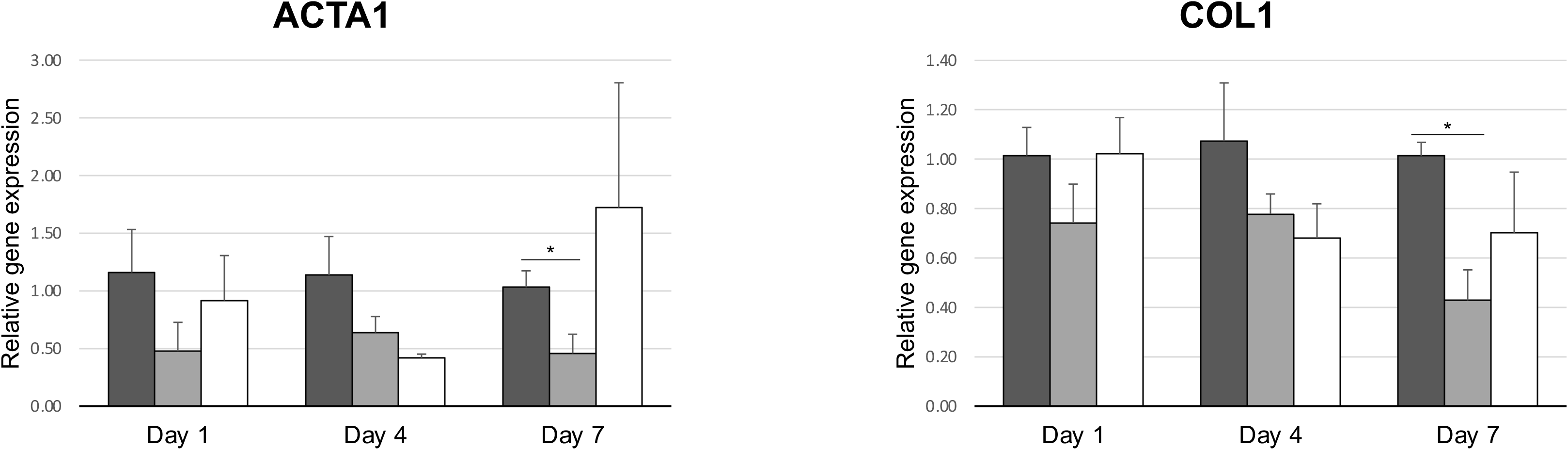
RT-PCR Markers for Fibrosis. Fibrosis markers including ACTA1, ACTA2, ACTRAP, SMAD2, TGFB1 and COL1. n=4 for all groups. Statistical analysis using two-tailed t test, * P<0.05.

Furthermore, the gene expressions of the angiotensin II receptor-associated protein (ACTRAP) and SMAD2 were significantly reduced in 35 dyne/cm^2^ compared with 6 dyne/cm^2^ group at day 4. Moreover, we also found that the gene expressions of alpha-smooth muscle actin (ACTA2), and Col1 decreased in 15 dyne/cm^2^ compared to 6 dyne/cm^2^ on day 7. In addition, another fibrosis marker transforming growth factor beta 1 (TGFB1) did not show significant differences among the groups.

### Effect of Vimentin as a Function of Shear Stress Magnitude for Porcine Fibroblasts

To investigate the porcine fibroblasts under varying conditioned medium treatment, vimentin was chosen as the reflecting the fibrosis marker, preserving the cellular integrity and providing resistance against stress. The porcine fibroblasts were cultured for 4 days, and then, DAPI and vimentin-GFP were used for staining. In Fig. 4. A & B, we used ImageJ to test the intensity of the vimentin and utilized the intensity of vimentin minus the background as protein expression. From Fig. 4. B, the 35 dyne/cm^2^ had significantly increased the vimentin expression compared to the 0 and 6 dyne/cm^2^.

**Figure 4.**
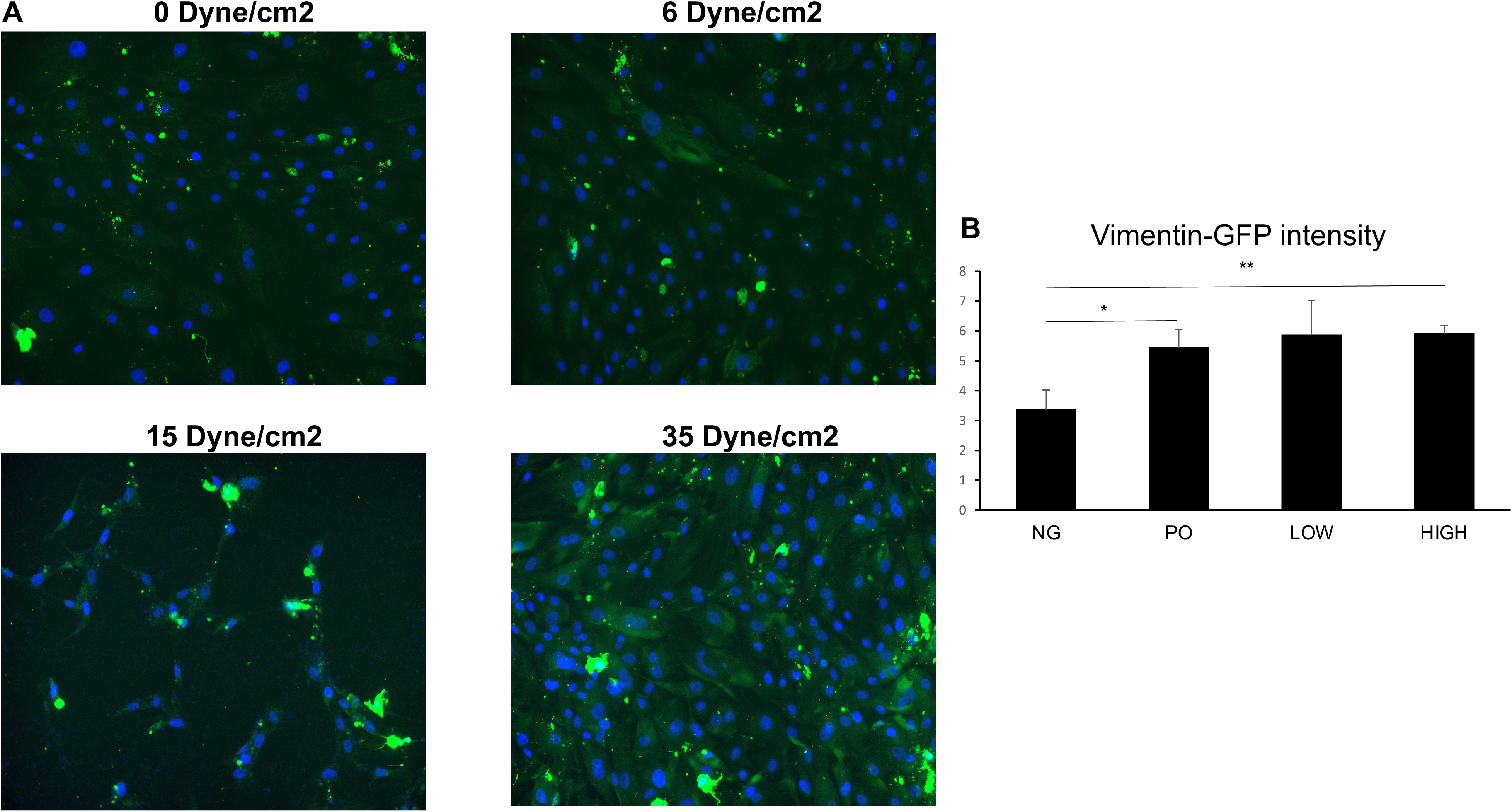
Vimentin Staining as a Function of Shear Stress Magnitude. Immunofluorescence of porcine fibroblasts at day 4 and day 7 under 25% conditioned medium. Vimentin was used as a marker for fibrosis. The intensity of Vimentin and the image background were analyzed by ImagJ. Data are expressed as mean±SEM. n=4 for each time point. Statistical analysis using two-tailed t-test, * P<0.05 and ** P<0.01.

### RNA-sequencing Revealed that Porcine Fibroblasts Induced Profibrotic and Proinflammatory Response

To explore the porcine fibroblasts metabolic pathway under conditioning medium treatment, RNA-sequencing was conducted to investigate the signaling pathways. From Fig. 5. A, the differentially expressed genes (DEG) demonstrated that the negative control had several genes modulated compared with the other groups, especially the high shear stress group including up and down-regulation. Typically, the negative control versus high shear stress group had a more significant increase than the other groups and reached at 1508 containing 647 down-regulated genes and 861 up-regulated genes. The low shear stress VS negative control and positive control vs negative control were also significant compared to other comparison groups. In Fig. 5. B, the volcano Plot histogram illustrates the relationship between statistical significance and the magnitude of variation. In the volcano Plot histogram, upregulated genes were situated on the right side, while downregulated genes were located on the left side. The upper regions represent genes that exhibit statistical significance [26]. Indeed, the data indicated that the positive control, low shear stress and high shear stress had upregulated the gene expression compared to the negative control. The top 10 upregulated and top 10 dow regulated genes, based on significance, for all 6 comparison groups, are shown in Fig. 5. C. This data shows the role of ECM modeling in promoting the fibrotic response of porcine fibroblasts during culture in EEC shear stress conditioned media.

**Figure 5.**
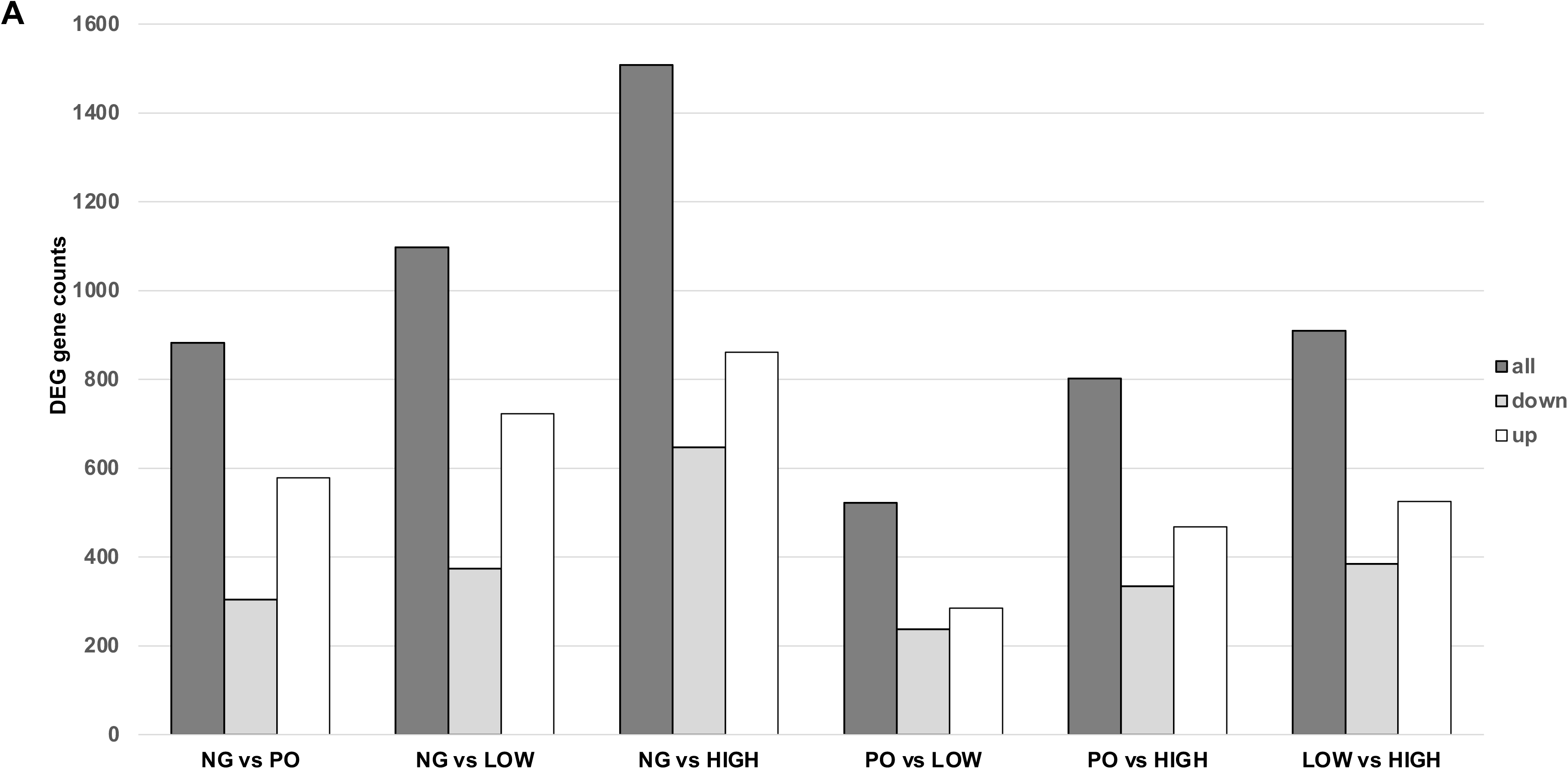
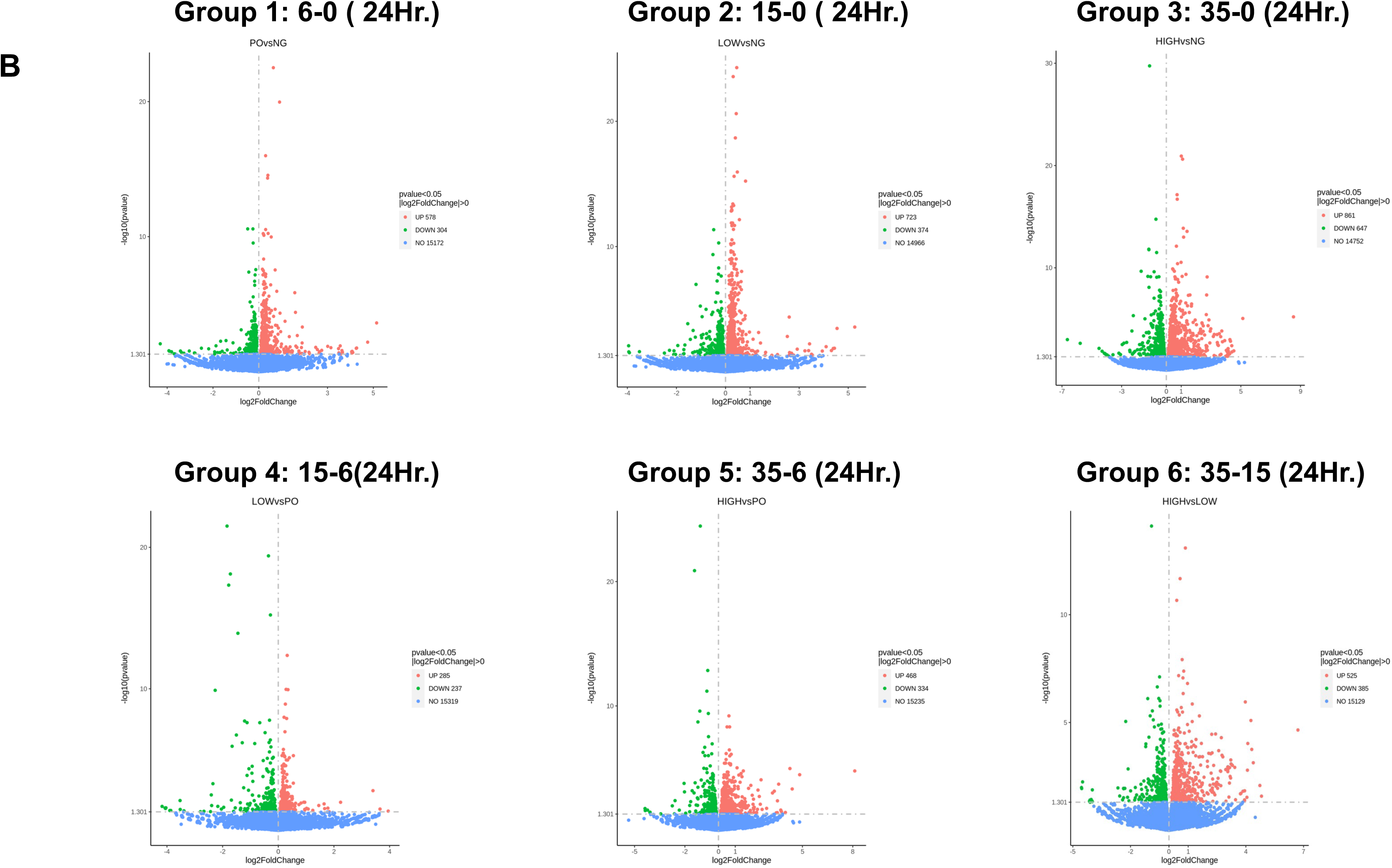
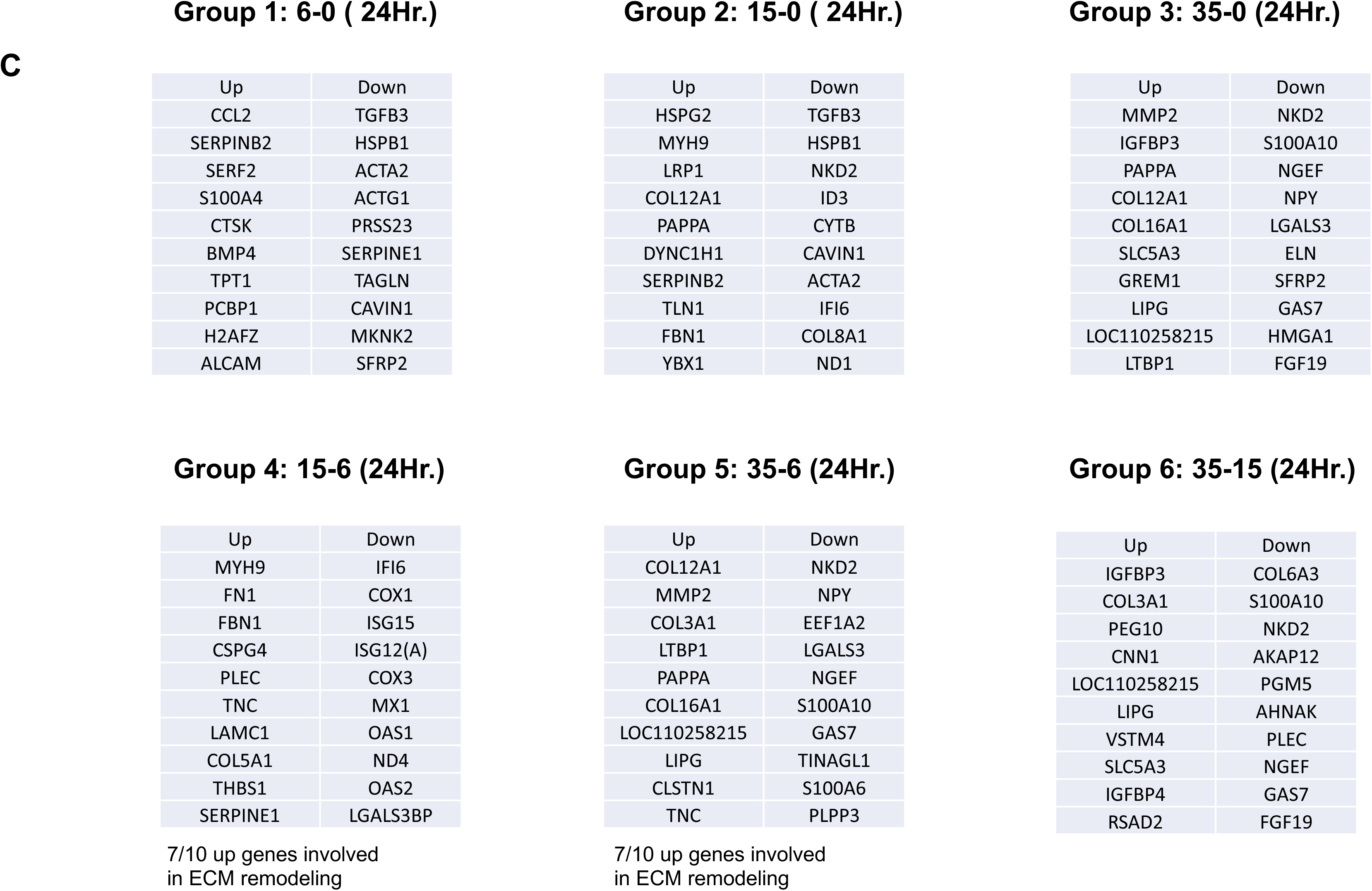
Differentially Expressed Genes. The statistics of the number of differential genes (including up-regulation and down-regulation) for each compare group and the threshold for screening are shown in a bar graph (A). The volcano plots are shown in (B) and all means the total number of differential genes in the compare group, up and down defined as the up & down-regulation number of differentially expressed genes in the compare group. The x-axis shows the fold change in gene expression between different samples, and the y-axis shows the statistical significance of the differences. Red dots represent up-regulation genes and green dots represent down-regulation genes. The top 10 upregulated and downregulated genes based on significane for all 6 groups are listed as individual tables.

The Fig. 6 was the images of the Gene Ontology Team which also defined as GO enrichment analysis. 30 Terms were categorized into major groups, including biological processes (BP), cellular components (CC), and molecular functions (MF). On the X-axis, there was the ratio of the gene linked to a specific GO term relative to the total number of differential genes. The Y-axis represents the GO terms, and the size of each dot corresponds to the genes annotated to GO term [27]. In Fig. 6, the negative control and positive control groups comparing with low and high shear stress groups showed that the membrane proteins, cytokines, cytoplasm, and organelles had a typically down regulation in the CC group. Moreover, the negative control and positive control groups comparing with low and high shear stress groups indicated that several pathways were increasing or up regulation of molecules in the MF group. Additionally, it shows that the high shear speed influences more up-regulation in the MF group from the data about the high shear stress VS negative control & low shear stress. The top 10 upregulated and downregulated pathways, based on significance are presented as a table for each of the six groups.

**Figure 6.**
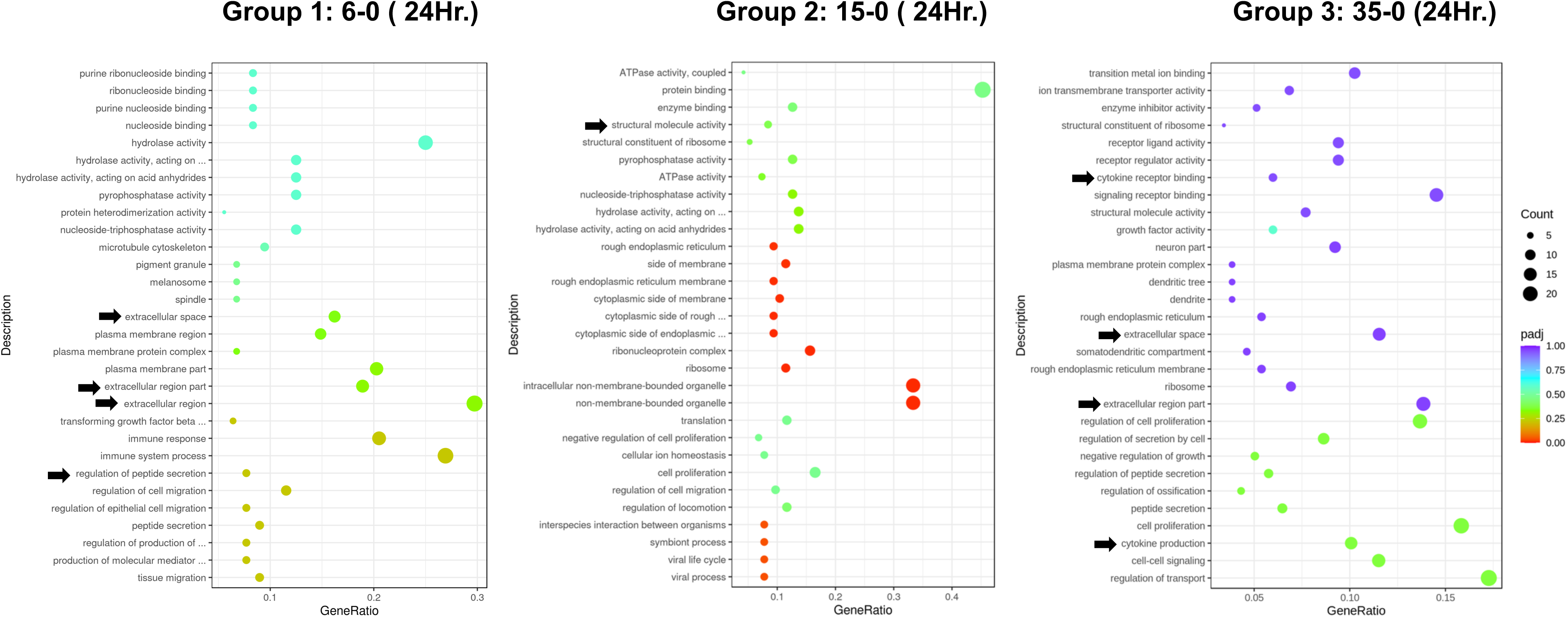
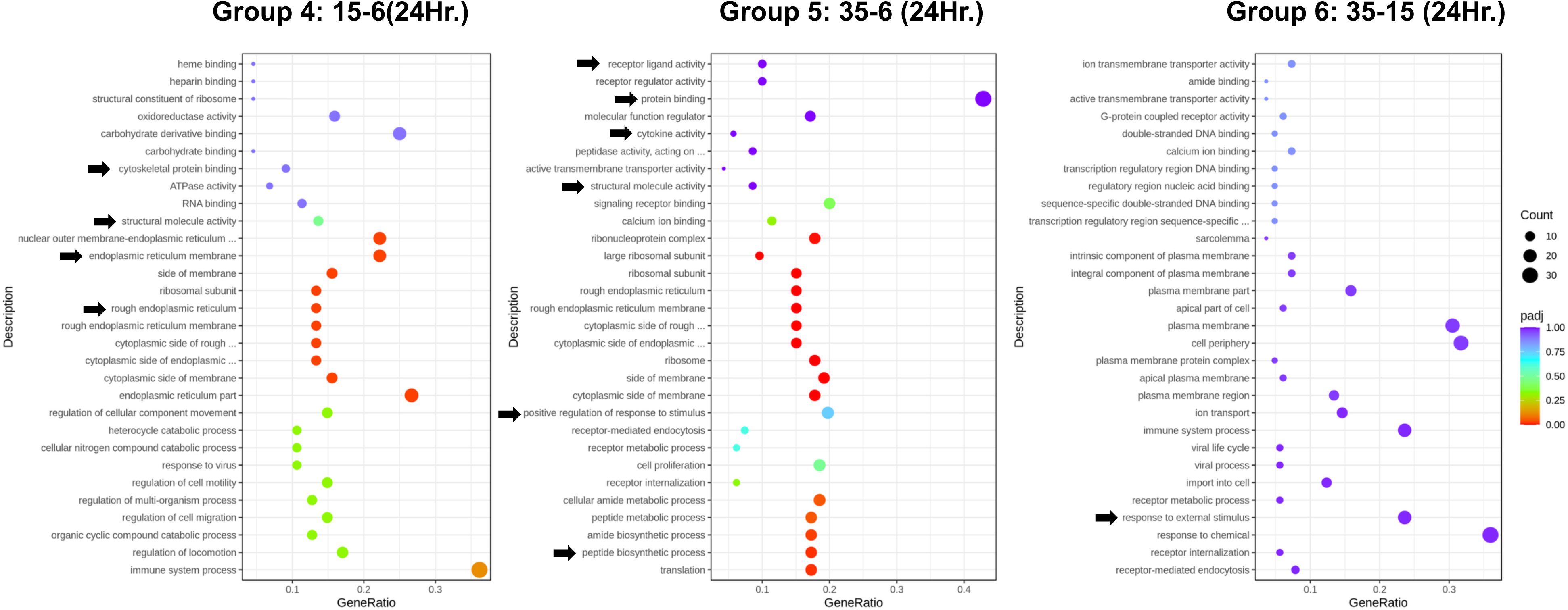
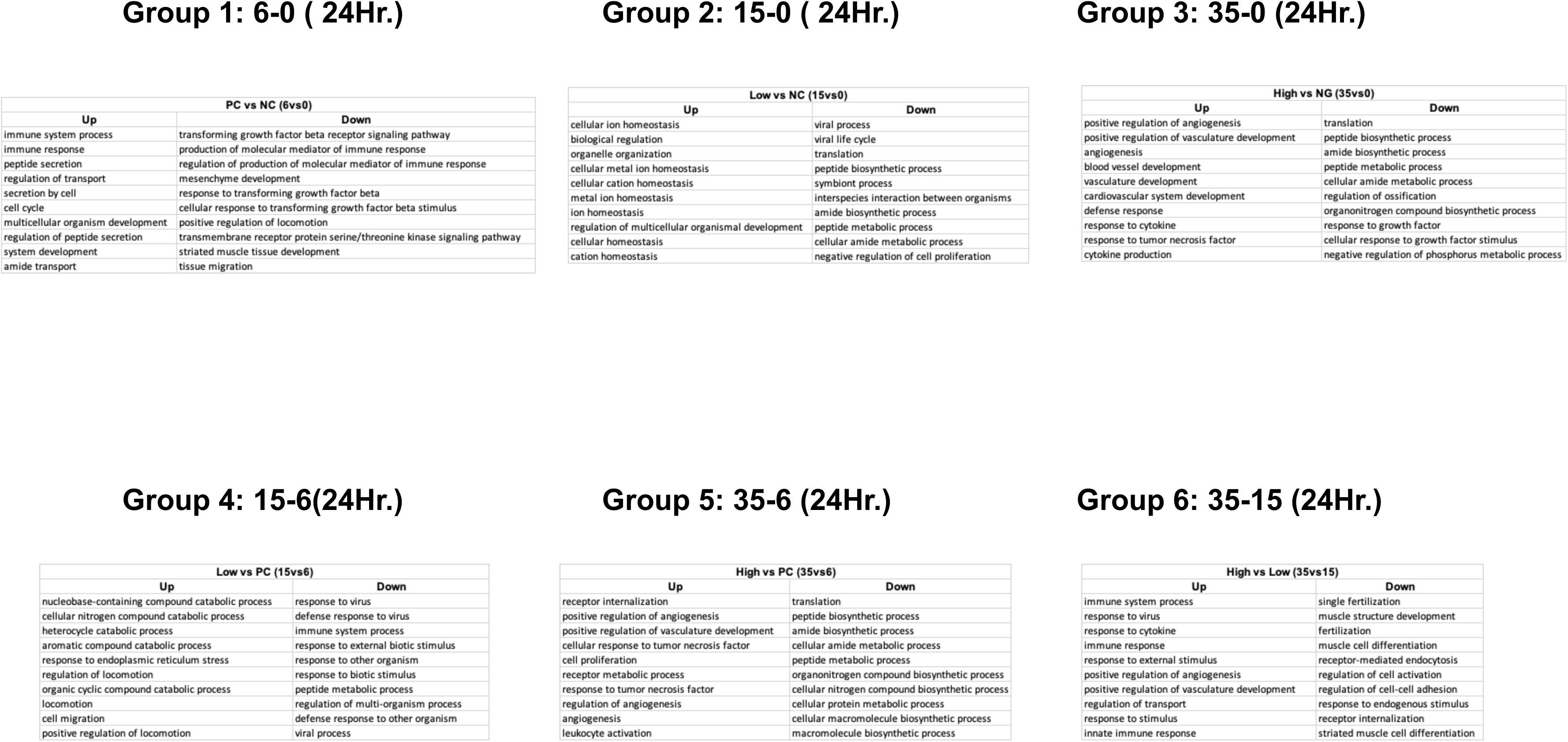
Gene Ontology Analysis. BP, MF, and CC represent Biological Process, Molecular Function, and Cellular Component groups of GO. The x-axis represents the ratio of differential genes linked with a given GO Term to the total number of differential genes, while the y-axis represents the respective GO Terms. The size of a point represents the number of genes annotated to a specific GO Term, and the color from red to purple represents the significant level of enrichment.

In the Fig.7 Kyoto Encyclopedia of Genes and Genomes (KEGG), functional assignment involves connecting the genes within the genome to the network of interacting molecules in the cell containing the metabolic pathways or cell signaling. Pathway enrichment analysis discerns significantly enriched metabolic, or signal transduction pathways linked to differentially expressed genes with comparing the entire genome [28]. From Fig. 8, the data demonstrated the potential significance of the inflammatory and fibrosis signaling pathway, including TNF-α, NF-κB, HIF-1, IL-17, PI3K-AKT, MAPK, etc., and the stimulation of immune cell responses. The top 10 upregulated and downregulated pathways based on significance are presented for each of the 6 groups. The most significant 11 enriched pathways are presented in Figure 8 and clearly demonstrate the profibrotic and proinflammatory response of the porcine firboblasts in response to porcine EEC conditioned media.

**Fig. 7.**
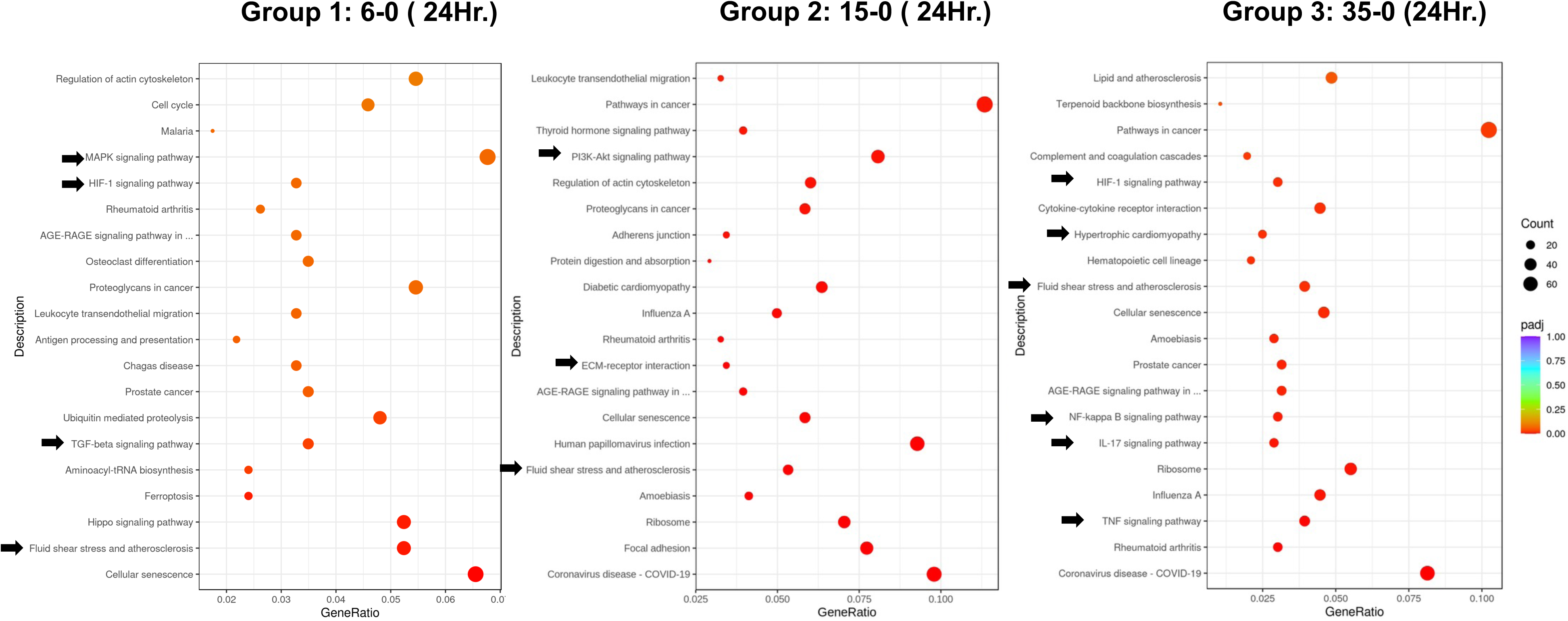
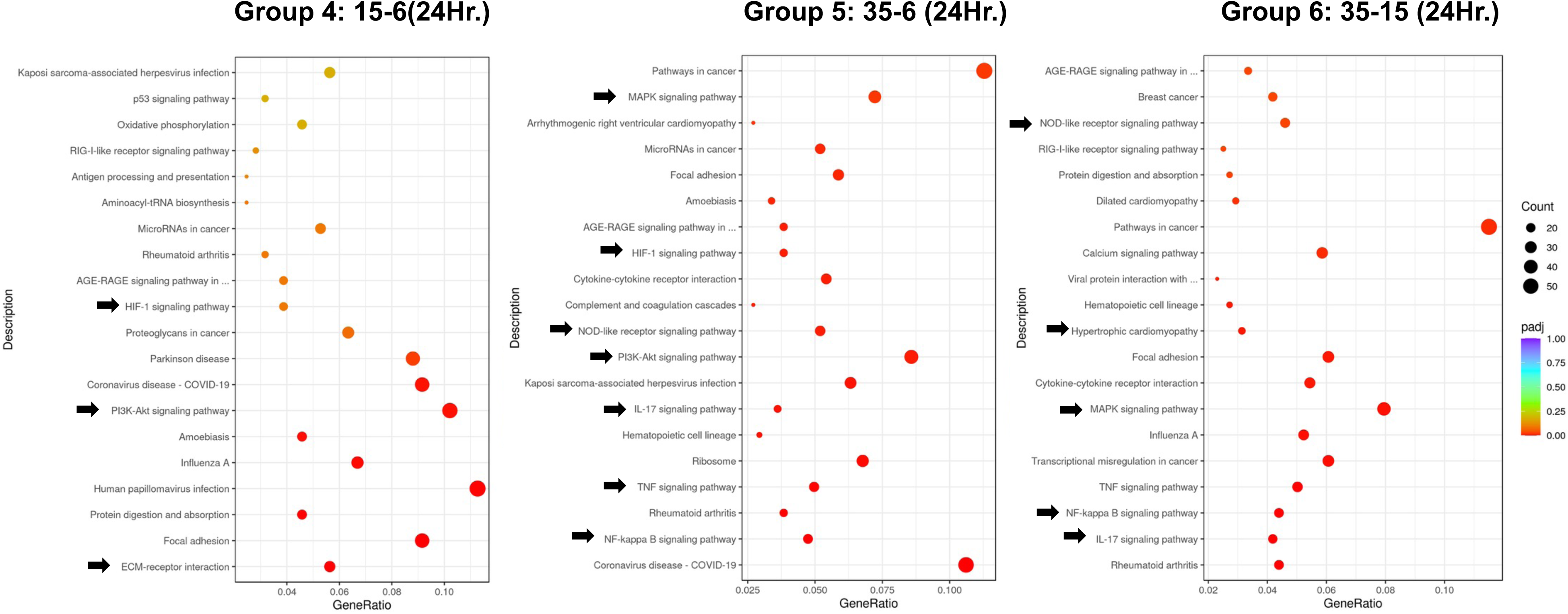
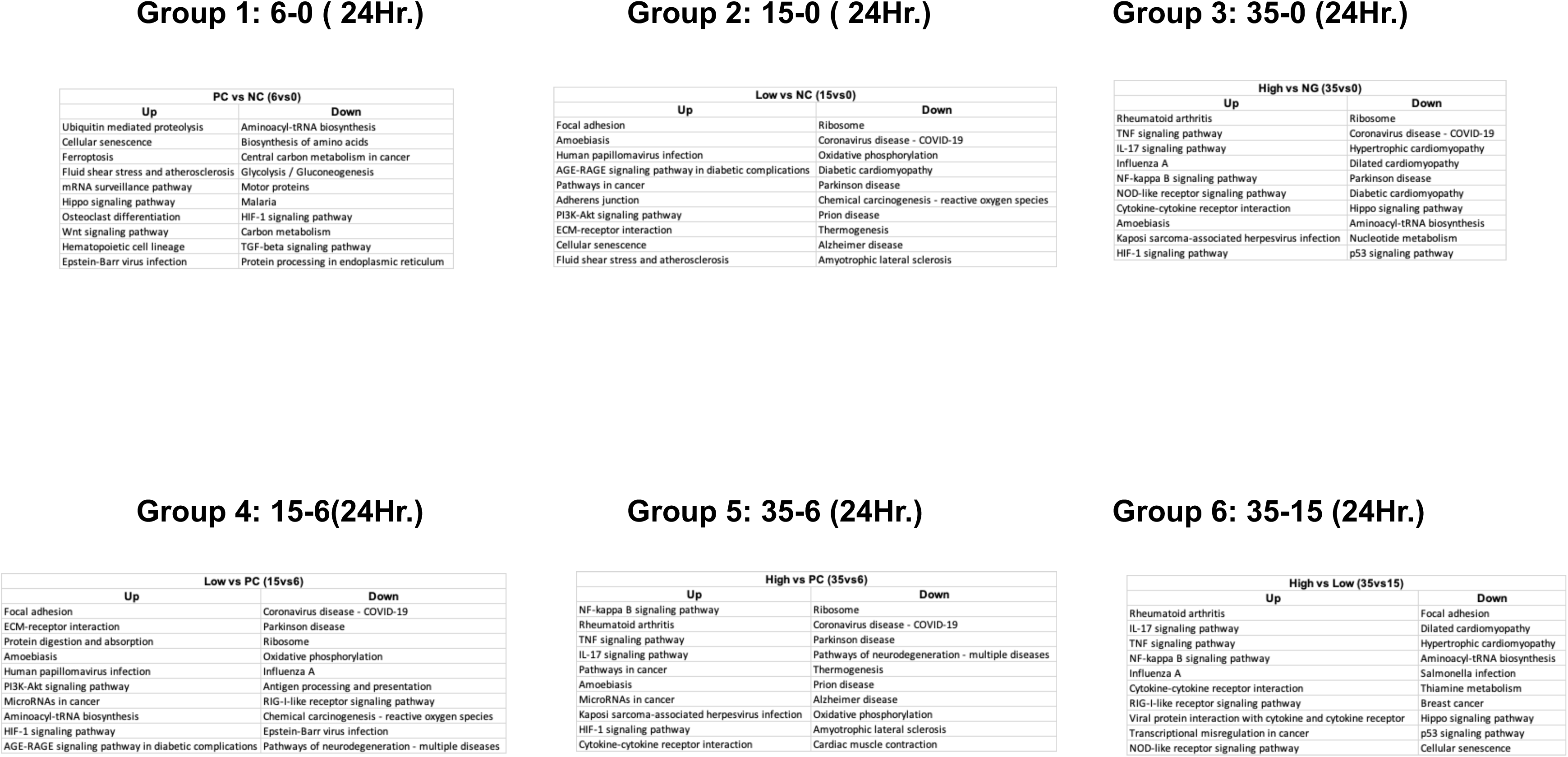
KEGG Pathway Analysis. The abscissa in the graph is the ratio of the number of differential genes on the KEGG pathway to the total number of differential genes, and the ordinate is KEGG pathway. The KEGG node including up-regulated genes is marked red, and the KEGG node including down-regulated genes is marked green. While yellow means the node contains both types of genes. Hovering over the marked KEGG node will show details of differentially expressed genes, with the same color as above, and the number inside brackets is log2.

**Fig. 8.**
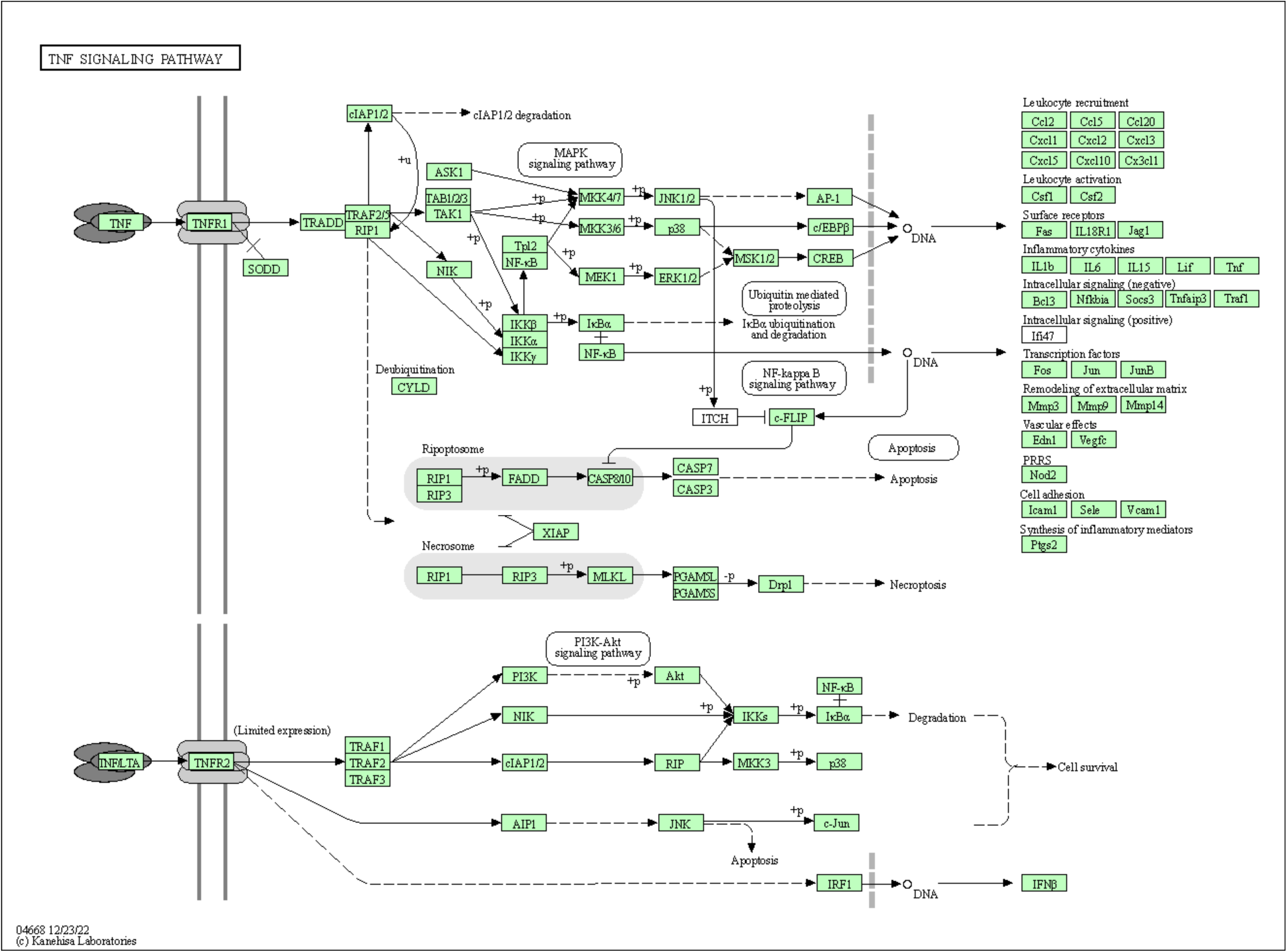
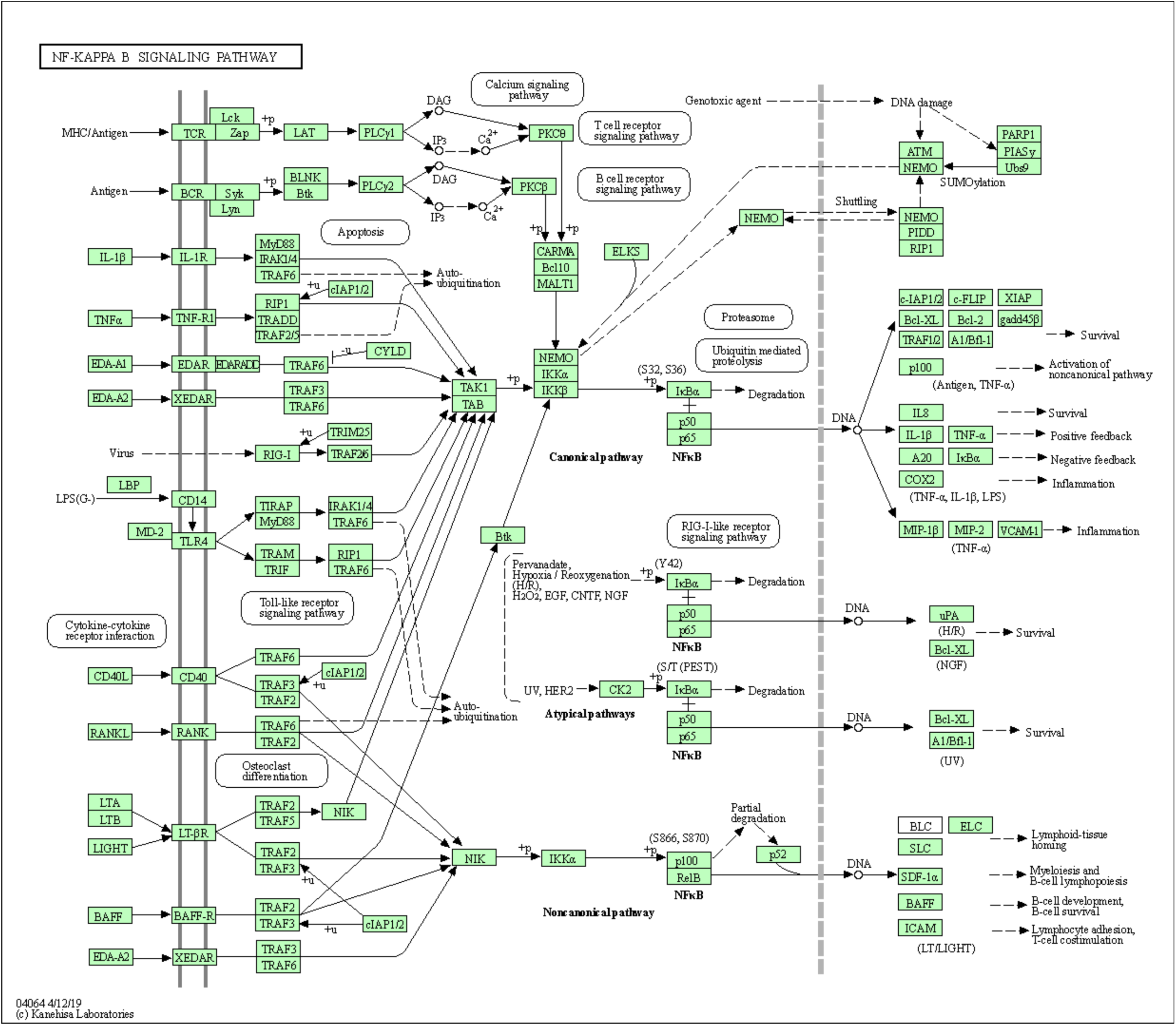
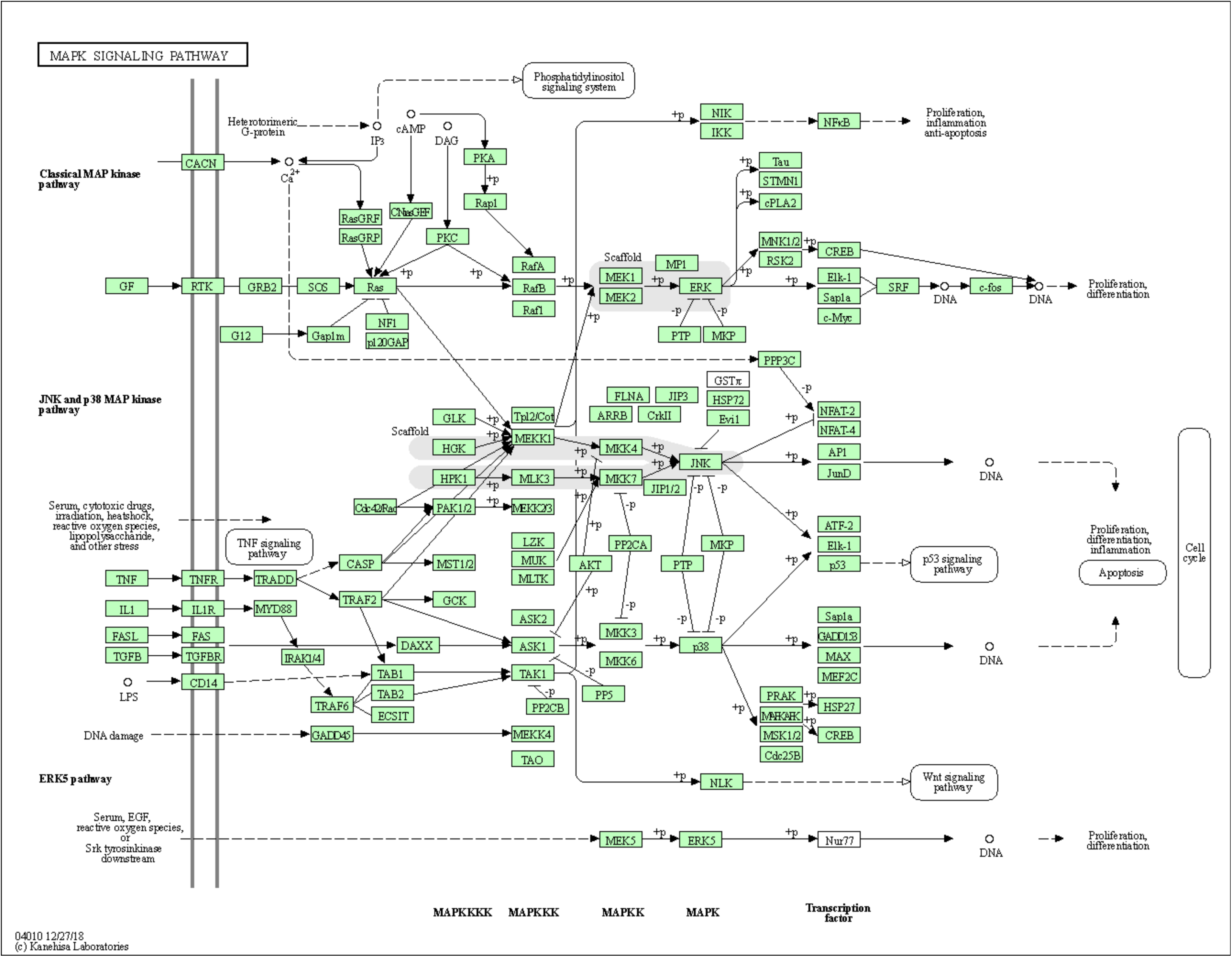
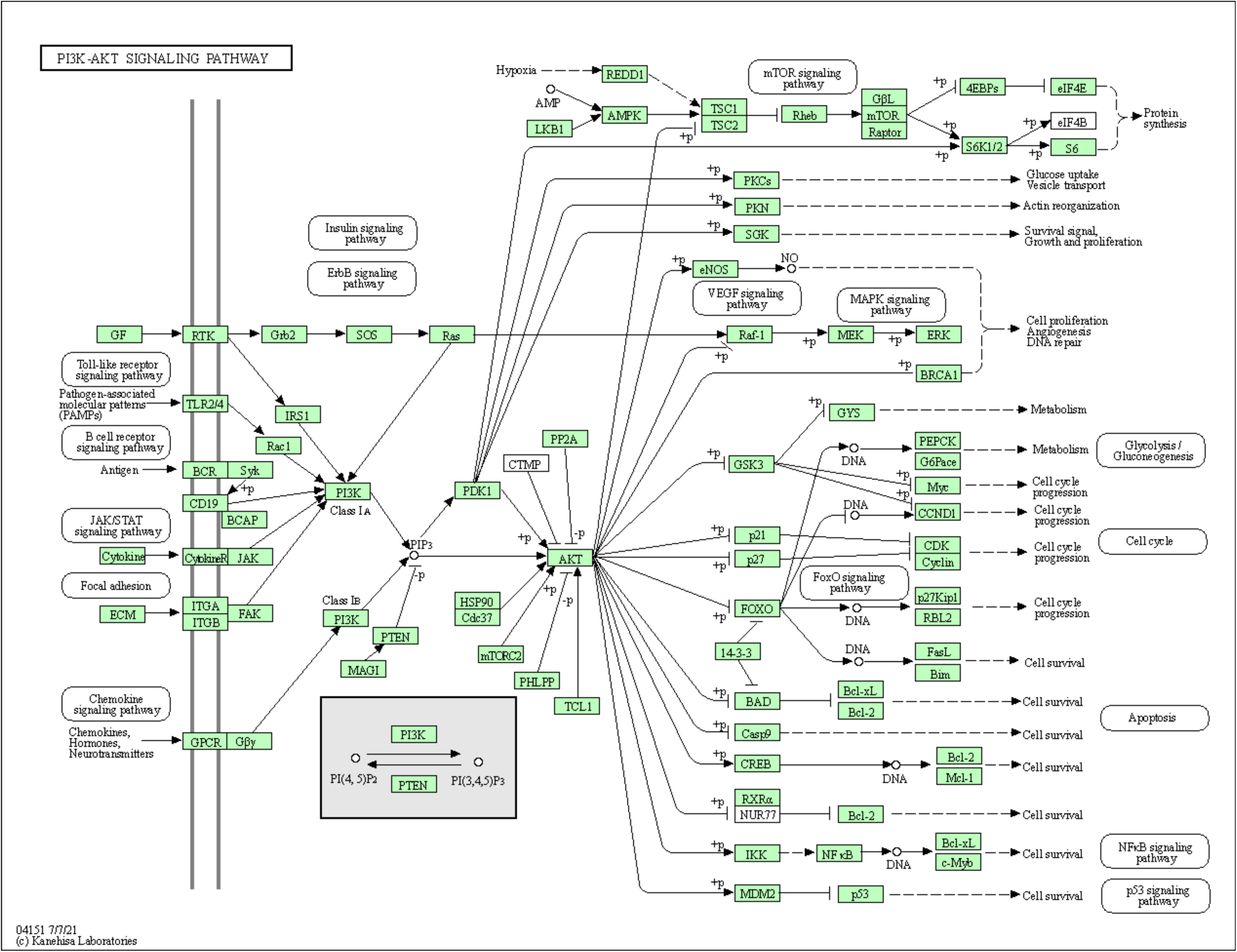
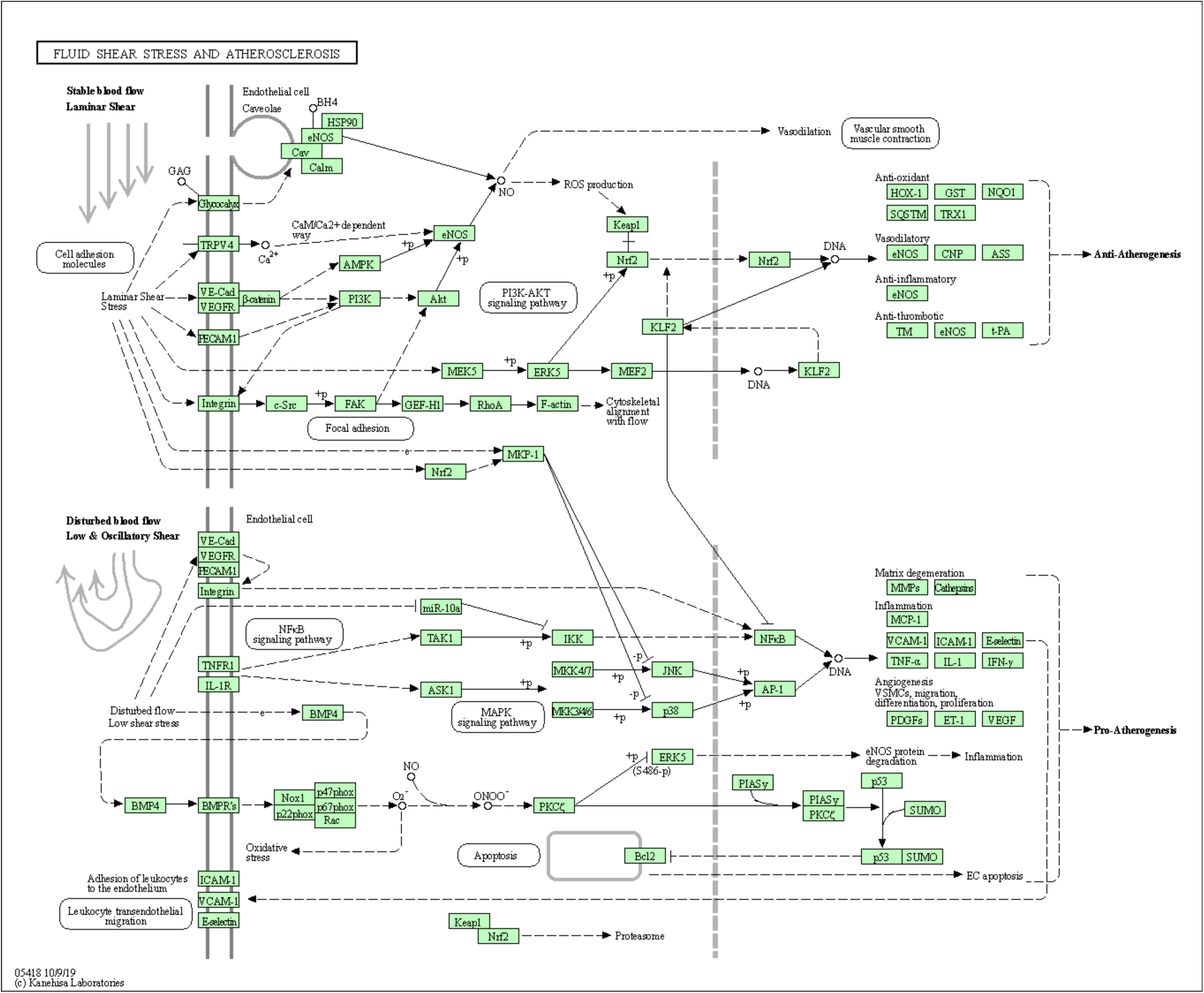
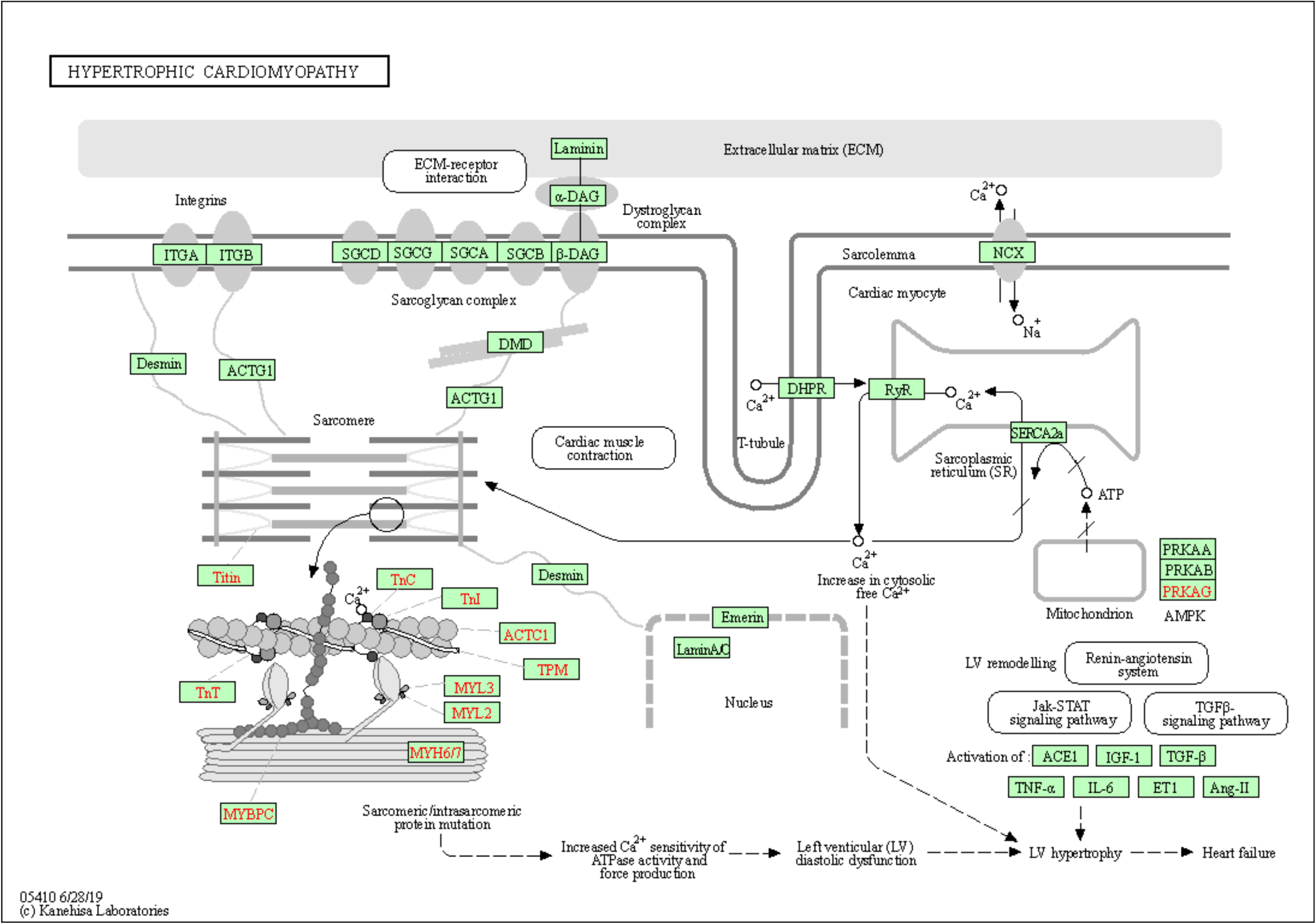
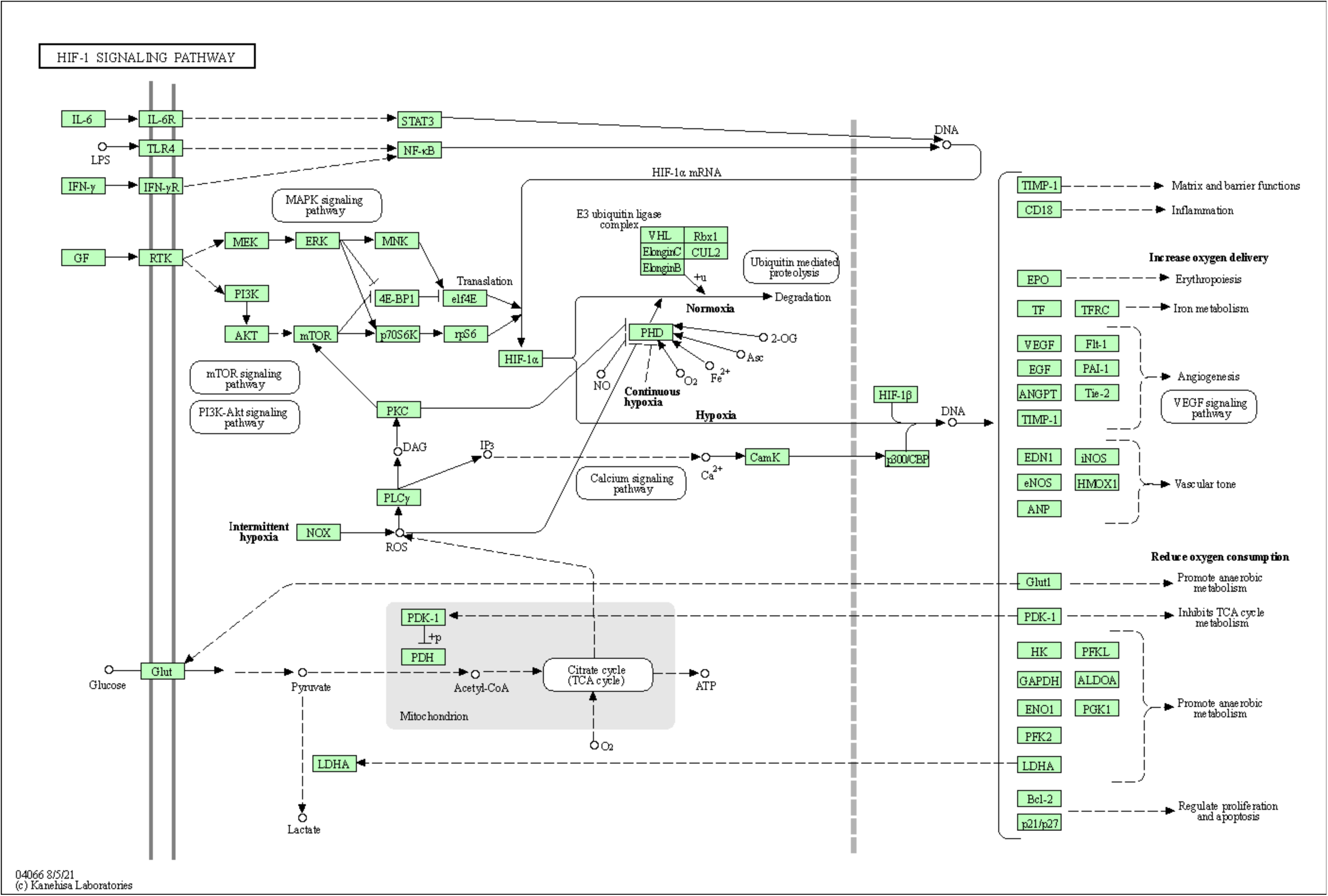
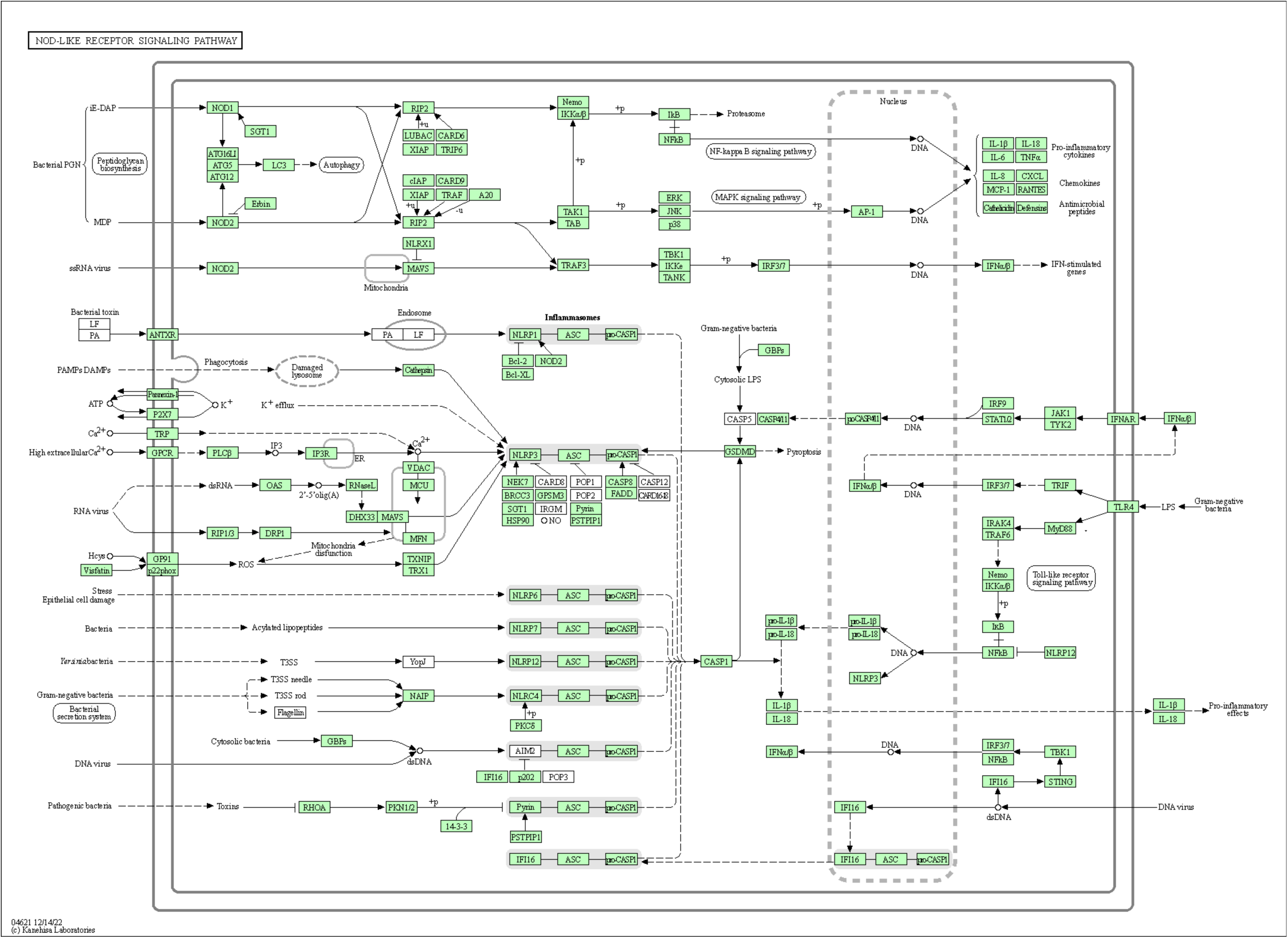
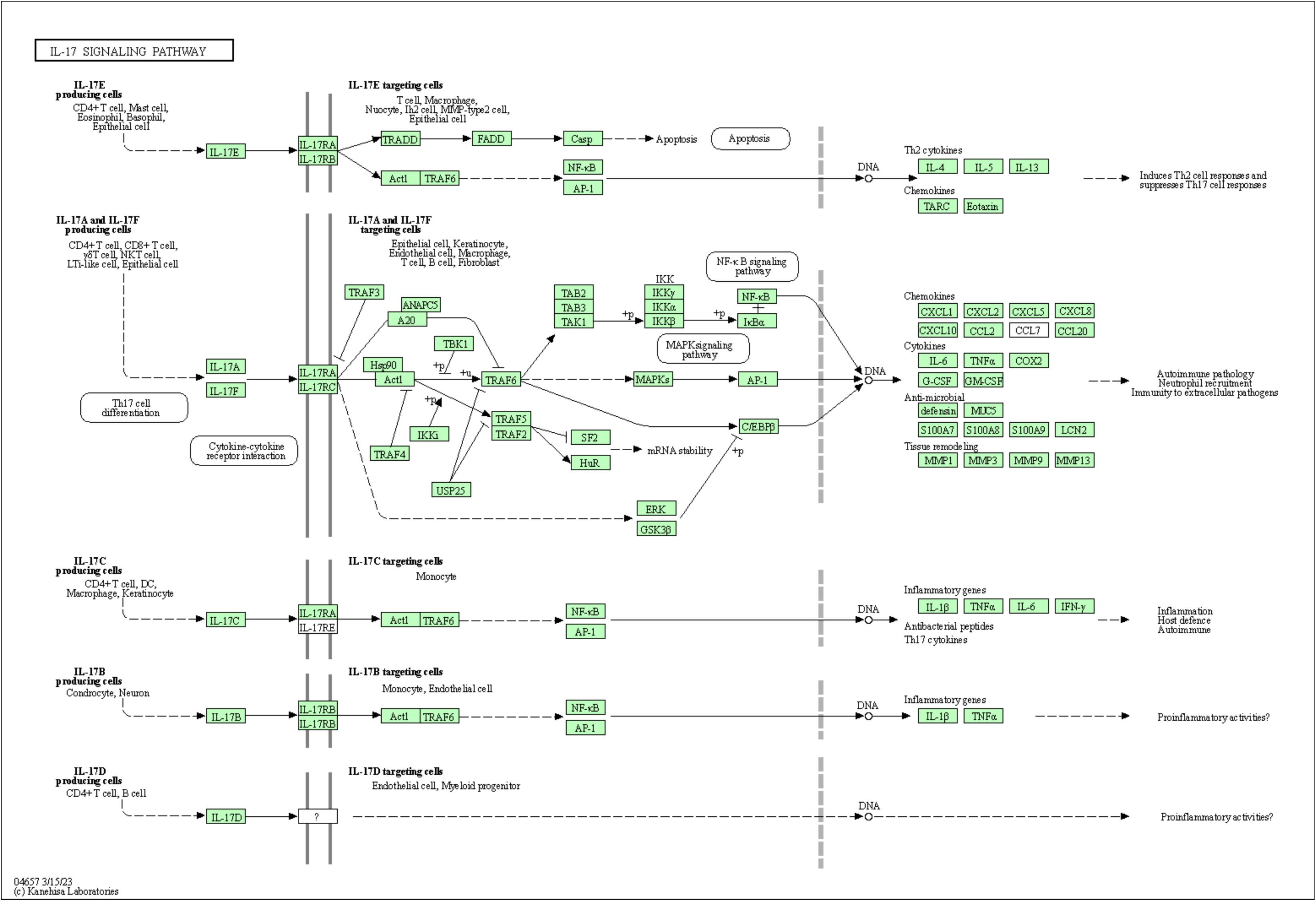
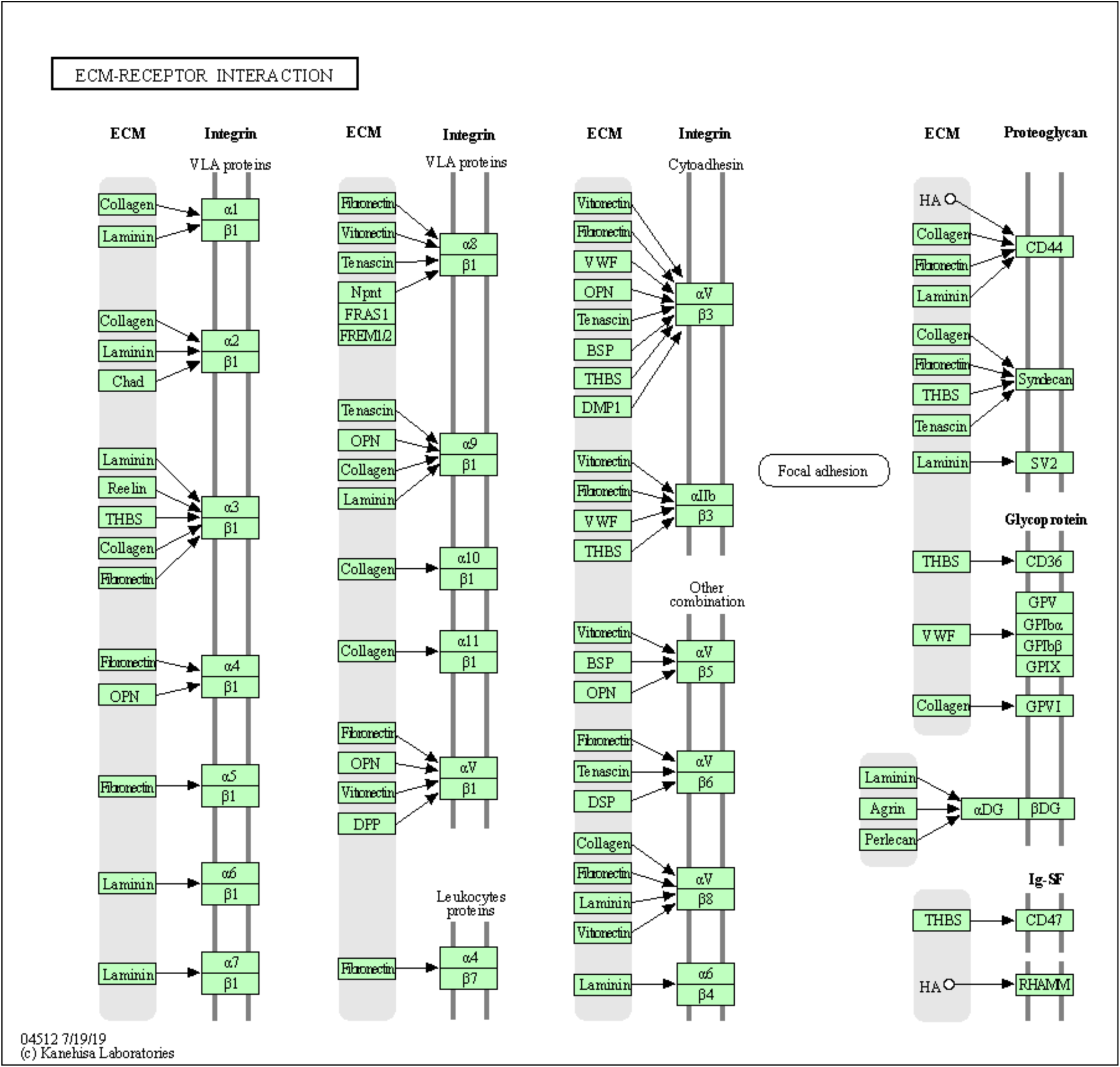
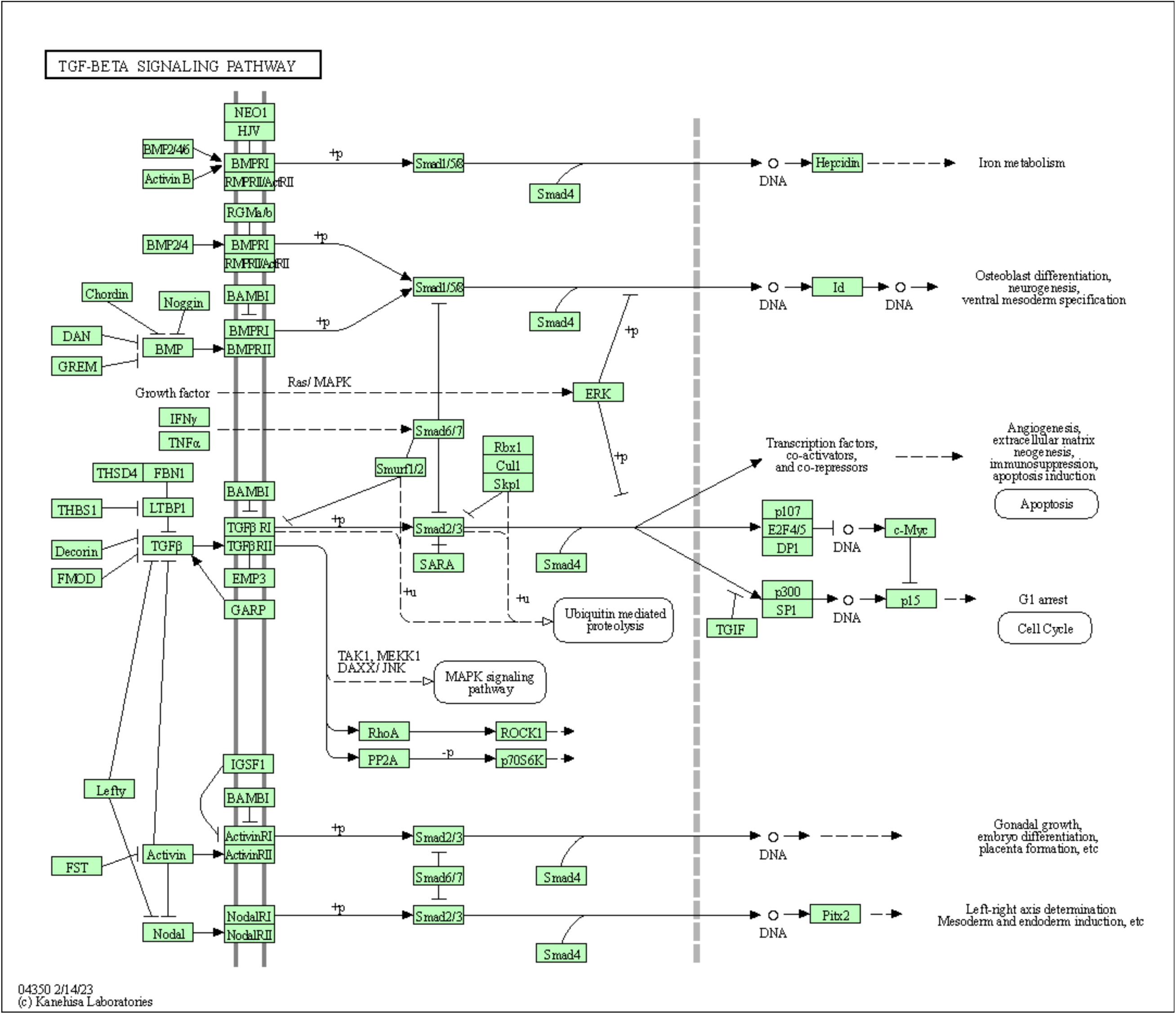
The most enriched pathways Based on Significance.

In summary, the RNA-sequencing data revealed that bioreactor treatment could induce inflammatory cytokines, and fibrosis and elicit an immune response in porcine fibroblasts.

## DISCUSSION

Discrete subaortic stenosis is a serious pediatric cardiac condition that is characterized by the formation of a fibrotic membrane in the left ventricular track outflow track (LVOT). We hypothesized the changes in the shear stress environment within the LVOT in DSS patients act on endothelial endocardial cells which release proinflammatory and profibrotic cytokines into the local environment. Subsequently, these proinflammatory and profibrotic cytokines act on the underlying fibroblast to promote the formation of the fibrous membrane. To test this hypothesis, in an earlier study, we quantified the magnitude of fluid shear stress within the LVOT of DSS patients and used cone and plate bioreactors to show the proinflammatory response of endocardial endothelial cells (unpublished data). The current study is a follow-up to this earlier work and the second step in the signaling cascade to investigate the effect of proinflammatory cytokines released by endocardial endothelial cells in response to shear stress.

The first step in our study was to optimize the ratio of conditioned media that can be used to culture porcine fibroblasts. We tested the effect of 10%, 25%, and 50% conditioned media on the viability of porcine fibroblasts. The high ratio (50%) of the conditioning medium with high shear stress (35 dyne/cm^2^) treatment had some cytokines that may cause fibroblast death, and it might be the inflammatory cytokines or could also be something else in the conditioned media. In addition, it appears that the normal porcine EEC medium without bioreactor treatment did not exert a detrimental effect on the proliferation and growth of the porcine fibroblasts in the negative control group when compared to other groups. However, in the high ratio of conditioned media group, the positive control group also decreased the cell number maybe due to the high concentration of conditioned medium, which may also include the high concentration of cytokines that caused the decreasing of the cell number of the porcine fibroblasts. The medium may have the pro-inflammatory cytokines and trigger the fibroblasts’ protection against the immune system and become fibrosis. The porcine EEC conditioning medium with low and high shear stress results in porcine fibroblast cell death, while the normal porcine EEC medium did not induce mortality in porcine fibroblasts. With increasing the shear stress magnitude and subsequently the rotational speed of the cone and plat bioreactor and high ratio of conditioned medium, which can cause more porcine fibroblasts to die. Therefore, from the results, the 25% porcine EEC conditioned medium was the suitable choice for this following research.

The RT-qPCR was to investigate the fibrotic response of porcine fibroblasts in response to treatment with conditioned media. ACTA1, ACTA2, COL1, TGFb1, SMAD2, ACTRAP were chosen as the representative fibrosis markers. ACTA1 is a member of the actin protein family, which plays a crucial role in cellular movement and the contraction of muscle fibers. Mutations in the ACTA1 gene are associated with the development of interconnected congenital myopathies [29]. From the ACTA1 data, the low shear stress group decreased the gene expression compared to the positive control group. A decrease in ACTA1 expression in fibrosis may be due to muscle dysfunction. In the context of conditions associated with fibrosis, this could manifest as muscle weakness or dysfunction [30]. Fibrosis involves an excess of tissue scarring and stiffening caused by the deposition of collagen. Reduced levels of ACTA1 might be linked to modifications in the structure and function of muscle tissue, potentially contributing to fibrotic changes [31].

Also, a decline in ACTA2 expression as well as the ACTA1 group may indicate a reduction in both the population and activity of myofibroblasts. It has the potential to diminish the deposition of collagen and other extracellular matrix components typically associated with fibrosis [32].

Reducing the expression of AGTRAP may serve as a therapeutic strategy to alleviate cardiac damage associated with fibrosis. AGTRAP paly a crucial role in the Angiotensin II (AII) pathway, and it is a strategy widely used in the treatment of conditions such as congestive heart failure [33]. A decrease in SMAD2 in the context of fibrosis implies a potential regulation alteration in TGF-β signaling pathway. SMAD2 serves as a crucial mediator in this pathway, which means that a decline in SMAD2 could potentially attenuate or restrict fibrosis by disrupting TGF-β signaling pathway [34]. From the above, we discovered the fibrosis marker alternations in the research, and the data indicated that the porcine fibroblast was reducing the gene expression of fibrosis markers due to the protection against to become further fibrosis from the conditioned medium stimulation.

Vimentin has been widely utilized to label cardiac fibroblasts [35]. Increased vimentin expression was observed around myocardial cells undergoing remodeling [36]. Vimentin serves as an indicator of inflammation and fibrosis through the regulated activation of the NLRP3 inflammasome [37]. Our results showed that the porcine fibroblasts had increased vimentin expression by IHC staining, and the high shear stress group dramatically improved the expression compared to other groups, which means that the shear stress induced the porcine fibroblast became fibrosis and inflammation.

Furthermore, we choose bulk RNA-sequencing from Novogene to investigate the gene expression of the porcine fibroblasts. From the results in the figures, we found that the NOD-like receptor, TNF-α, NF-κB, HIF-1, IL-17, PI3K-AKT and MAPK signaling pathways were activated or upregulated in the high shear stress group compared to other groups. It has been reported that the TNF-α promotes the stabilization of snail and β-catenin proteins by hindering GSK-3β-mediated phosphorylation through the NF-κB and AKT signaling pathways [38]. The MAP kinase signaling pathways trigger a secondary response by upregulating the expression of various inflammatory cytokines, including TNFα, thereby enhancing the biological activity of TNFα. Consequently, MAPK operates both upstream and downstream of TNFα receptor signaling [39]. The IL-17 signaling pathway commences the signaling through Act1-induced K63-linked ubiquitylation of TRAF6, thereby activating the MAPK pathway, CCAAT-enhancer-binding protein β (C/EBPβ), and NF-κB pathways [40]. Some literatures demonstrated that HIF-1α signaling has been associated with the development of fibrotic diseases affecting diverse organs. But the specific activation and functional influences of HIF-1α signaling in fibroblasts during the progression of fibrogenesis remain incompletely understood [41]. Therefore, the HIF-1α signaling pathway may also indicate that the porcine fibroblast became fibrosis under the conditioned medium treatment. Apart from that, accumulated studies implicated that inhibiting the PI3K/Akt pathway may alleviate inflammation and fibrosis induced by BLM, potentially reversing the lung fibrosis-associated epithelial-mesenchymal transition (EMT) process [42]. The activity of p38 MAPK has been strongly linked to the development and progression of diseases in inflammation and fibrosis [43]. Based on the literature, p38 MAPK has emerged as a therapeutic target for chronic inflammatory and fibrotic conditions. Recent studies have reported that pharmacological inhibition of p38 MAPK effectively mitigated cardiac fibrosis in a mouse model [44]. Additionally, it has been demonstrated that a p38 MAPK inhibitor prevented myofiber damage and muscular fibrosis in a mouse model of muscular dystrophy [45].

To summarize, the discussed signaling pathways are intricately connected to inflammation and fibrosis. These data are associated with the observed phenotype resulting from the treatment of fibroblasts with condition medium. The RT-qPCR and immunofluorescence data exhibit significant concordance with the RNA-seq data, showing consistent patterns and alignment in multiple aspects. Specifically, activation of fibrotic markers and inflammation was evident under treatment with high shear stress conditioned medium. Therefore, our results serve to support our hypothesis that cytokines released from endocardial cardiac cells in response to shear stress response in the LVOT are responsible for driving the profibrotic response in fibroblasts and subsequently, formation of the fibrotic tissue in DSS patients. These studies provide a pathway to develop a novel therapeutic strategy to treat this vulnerable patient population.

## ACKNOWLEDGMENTS

This work with supported by an NIH R01 Grant number 5R01HL140305. This work was supported by funding from the Lew and Laura Moorman Family Foundation to SGK. The authors would like to thank financial support from the Division of Congenital Heart Surgery, Texas Children’s Hospital, Houston, TX.

## Notes

### Competing Interest Statement

The authors have declared no competing interest.

